# Cluster-Based Inference for Memory-Based Cognition

**DOI:** 10.1101/2022.04.22.489185

**Authors:** Will Penny, Nicho Menghi, Louis Renoult

## Abstract

This paper proposes a model of hippocampal-based category learning using a cluster-based inference framework which produces two systems (i) an extendable cluster-based memory module (CMM) that provides representations of learning episodes with strong pattern separation, and supports online decision making during that learning, (ii) cluster-based task modules (CTMs) which consolidate clusters learnt by CMM to support higher-accuracy decision making in the mid-term. Learning in CMMs optimises the joint probability of stimuli and category labels, whereas learning in CTMs optimises the probability of categories given stimuli. The latter system learns from the former via a process of “cluster consolidation”. We apply the model to data from a behavioral learning task and find that, as well as improving categorisation performance, cluster consolidation decreases recognition scores for old cues but increases them for novel cues. This model-based perspective explains forgetting and false memory effects as serving future categorisation performance. The paper also expresses a view that models of memory-based cognition should provide human-level performance on complex categorisation tasks, and do so with minimal labelled data. In working toward this goal we therefore compared the accuracy of CMM- and CTM-based decision making to standard Softmax Regression approaches on benchmark machine learning datasets. This produced mixed results. We found some significant improvements of CMM over Softmax Regression and of CTM over CMM. Overall, our framework reconciles complementary learning systems theory with more recent findings from cognitive neuroscience of generative replay and hippocampal-based generalisation.

## 1 Introduction

Complementary Learning Systems (CLS) theory [46] proposes that the mammalian brain possesses two learning systems with complementary properties. The first is anatomically based in hippocampus and quickly learns the specifics of individual experiences whereas the second is based in neocortex and gradually learns structured knowledge representations, thus facilitating generalisation. CLS was based on the connectionist assumption that neocortex uses representations of the sort you find in a multilayer neural network i.e. highly distributed, multi-layered and with units using non-local basis functions. The driving concern of CLS was to explain how neocortex could learn without suffering from catastrophic forgetting, whereby knowledge gained about one task is overwritten when learning another. The insight of CLS was that catastrophic forgetting could be avoided by interleaving samples of older tasks with samples of new tasks – using what is referred to in current terminology as “experience replay” [71].

The connectionist assumptions underlying CLS seem prescient given the recent success of neural networks in Artificial Intelligence (AI). The success of deep learning has shown that (augmented) connectionist architectures can provide human level learning across tasks such as image and speech recognition. They therefore seem fit for purpose from a functional perspective.Moreover, brain imaging has shown that the representations learnt by these networks provide the best explanation to date, for example, of feedforward hierarchical processing in the human ventral visual stream [72]. Further, the use of experience replay and interleaved sampling provides the basis for artificial multi-task and continual learning systems that provide state-of-the-art performance across a range of benchmark data [52, 67].

Two more recent strands of evidence from neuroscience research, however, presents a more nuanced picture of how the human brain might solve the continual learning problem. First, studies of replay activity are consistent with the idea of “generative replay” [71] in which the hippocampal system is thought to capture the statistical structure within exemplars and later draw samples from that statistical model. Second, in addition to its well-established function in episodic memory, the hippocampal system is also thought to be able to generalises across experience [66], a phenomenon referred to as “hippocampal-based generalisation” [73].

This paper presents a computational model that updates CLS to accommodate generative replay and hippocampal-based generalisation. We focus on hippocampal-based learning and the sorts of representations that would be useful for both teaching neocortex and providing interim decision making support until a neocortical system is ready to take over. Our model therefore also addresses the little attended question “if the neocortical system is slow to learn, how are decisions made in the interim?”. Since CLS was first proposed there have been additional proposals for how it may be updated in light of more recent findings [51, 42]. We return to this topic in the discussion.

This paper proposes the decompositon of hippocampal-based learning into the interaction of two systems: (i) a cluster-based memory module (CMM), which learns the statistical structure of experiences and can support online decision making during that learning and (ii) cluster-based task modules (CTMs) which create new representations based on those provided by CMM. This results in better decision making in the mid-term. Because both of these systems use cluster-based representations, creating one from the other is more straightforward than creating an entirely different neural-net based representation. We refer to this latter (neocortical) system as the set of neural-net-based task modules (NTMs) which support decision making in the longer term, after systems consolidation is complete [46]. NTMs reflect the functional specialisation of neocortex with networks sharing common lower-level processing layers but having different task-specific output layers (in machine learning this is known as a multi-headed structure, with a different “head” or output network for each task). Generally, we expect that optimal task performance is provided by NTMs, but there may be types of categorisation or reward learning problems for which this is not the case (ones in which the mapping from input to output space can be economically partitioned into a set of input regions in which the output does not vary, or varies smoothly – this is the type of representation used in CTM). Or it may be that full transfer of decision making to NTMs is only complete in the large-data limit. Tasks for which CTM can provide a high-level of performance will likely remain hippocampal-dependent for longer.

We illustrate the overall approach with the problem of category learning, as shown in Fig 1. Here, CMM clusters are optimised to maximise the joint density of (compressed) multisensory input and category labels, whereas in CTM, these clusters are merged so as to maximise the conditional density of category labels given multisensory input. We refer to this as “cluster-level consolidation” as opposed to the systems-level consolidation that results in the development of representations in NTMs. The twin goals of producing a memory system with both (i) good pattern separation between similar experiences and (ii) good generalisation over similar experiences, are inherently opposed. Here, we resolve this tension by having strong pattern separation in the CMM and strong generalisation in the CTM.

**Figure 1:**
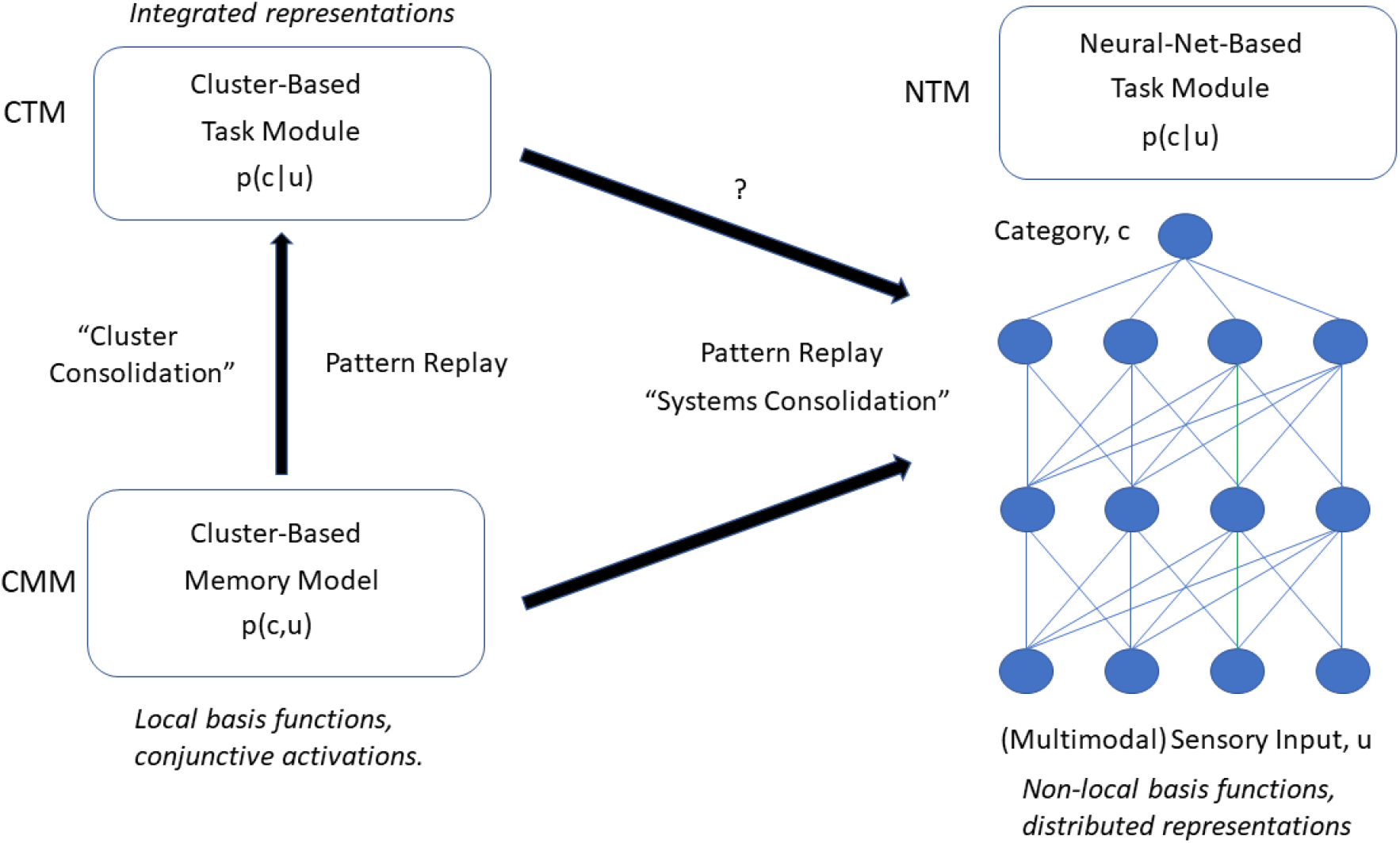
We consider hippocampal-based learning as being implemented in two systems (i) a cluster-based memory module (CMM) that learns the statistical structure of experiences and can support online decision making during that learning, (ii) cluster-based task modules (CTMs) which morph representations learnt by CMM to support higher-accuracy decision making in the mid term. This takes place via a process of cluster consolidation. Neocortical learning is instantiated using (iii) neural-net-based task modules (NTMs) which support gold-standard decision making in the longer term. In the context of a category learning task, CMM creates a joint density model *p*(*c, u*) over categories *c* and inputs *u*, whereas CTM and NTM create conditional density models of categories given inputs, *p*(*c|u*). This paper focusses on CMMs and CTMs.

We view the activity of these three learning systems as being consistent with complementary learning systems theory, transformational accounts of learning and memory [70], AI-based perspectives on brain function that emphasise the importance of system-specific cost-functions and architectures, and accounts of hippocampal-based generalisation from cognitive neuroscience. Ours is not the first paper to propose new perspectives on CLS – we relate our modelling approach to these other developments [51, 42] in the discussion.

### 1.1 Replay

The original CLS proposal [46] envisages the hippocampal system as receiving compressed multi-sensory input from, for example, neocortical autoencoder networks with the hippocampus itself then able to reinstate patterns that were experienced during behaviour at later offline periods, such as in sleep or quiet wakefulness. This reinstatement has been observed empirically in rats [22] and humans [66] and the broader phenomenon of “replay” has received much attention in the last decade – see Wittkuhn et al. [71] for a recent review of the various types of replay and proposed computational functions. For example, ‘interleaved replay”, in which exemplars from multiple categories or tasks are interleaved together, is proposed to support systems consolidation; ‘experience replay” refers to verbatim reinstantiation of experienced episodes as opposed to “model-based replay” or “generative replay” in which samples are produced from a model of previous experience. We note that a complementary learning systems perspective, making use of interleaved replay, is used in recent continual learning systems that provide state-of-the-art performance on benchmark machine learning problems [59, 50, 52, 67]. The CMMs propose in this paper are consistent with the idea of generative replay.

### 1.2 Cost Functions

Perspectives on the integration of neuroscience with recent advances in Deep Learning [18] advocate thinking about the brain in terms of the costs functions that may be optimised in different brain regions, and the structured architectures that may be required for dedicated systems such as attention, short- and long-term memory, and transfer learning.

So, what costs functions are implicit in CLS? In generative replay (see e.g. [63]), parameters of a generative model are estimated online so as to provide a model of the joint density of training data. For category learning, with input stimuli *u* and category labels c, this is the joint density *p*(*c,u*). During an offline period, e.g. sleep or waking rest, samples can then be generated from this model to provide data for neocortical learning. These samples will not be identical to the original data but will have the same statistical characteristics in so far as they are captured by the CMM. As we shall see below in the context of cluster-based models, original data can be replayed with high fidelity if there are a sufficient number of clusters each with a sufficiently high encoding precision. This correponds to the case of strong pattern separation. These samples can then be replayed to the CTM and NTM to learn the conditional probability of categories given stimuli, p(c|u).

In this paper we decompose the role of the hippocampus into two systems; the CMM and the CTMs. Learning in the CMM does not maximise the conditional density but creates a joint density model of categories and inputs (it can then predict categories from inputs using pattern completion – see below). However, it is well known in the statistics and machine learning literature [7] that discriminative models (based on the conditional density) provide more accurate decisions than generative models (based on the joint density). We therefore expect NTMs to be more accurate than the CMM – no surprise, because each NTM is a special purpose system to designed to implement a single task. The goal of the CMM is to capture and organise experience.

CTMs are also trained to maximise the conditional density, so will also be better categorisers than the CMM. However, we expect NTMs to be better categorisers than CTMs for the following reasons. First, the deep learning architectures that are thought to underly NTMs [72] provide state of the art categorisation performance – this will be better than CTM classifications that are effectively based on distance metrics (distances from exemplars to centres of clusters). Second, we consider learning in CTMs to be constrained to the search path provided by cluster merging. This produces fast learning but with sub-optimal performance. Generally, we would therefore expect NTMs to have better performance than CTMs (but see points above regarding the large-data limit and nature of the input-output mapping).

### 1.3 Architectures

What are the architectural features of CLS?

In the original CLS paper [46] the neocortical system was instantiated as a Multilayer Neural Network (MNN) and, indeed the two proposed properties of neocortical learning (that it is slow and provides generalisation) are consistent with MNNs and more recent deep learning approaches. Both properties derive from the highly distributed representations that the networks employ. These provide deep networks with their tremendous explanatory power whereby representations instantiated in shallower layers can be shared across tasks thereby facilitating knowledge transfer. This same property however makes learning slow – the optimal strength of a connection in a deeper layer depends on the values of connections in shallower layers, and vice-versa. These interdependencies make learning difficult, with the underlying optimisation problem containing multiple local maxima. In practice, effective learning can only take place “offline” i.e. with incremental parameter adjustments over repeated presentation of batches of training data [31].

In contrast, hippocampal representations are thought to be less distributed, less layered and to use more localised basis functions. CLS assumes a view of hippocampus as providing sparse coding such that unique experiences map onto the activation of a small number of active neurons. As we shall see, this is consistent with a cluster based coding where cluster activations are equivalent to localised basis functions (for example Gaussians). A good example of this are the hippocampal place cells. In this (simplified) view, hippocampal models also have two key features (i) processing units use local basis functions, and (ii) these units are combined into shallow networks. Learning is straightforward with such local representations as only a small numbers of parameters need updating for each exemplar presented. This is not the case for the distributed representations in MNNs.

### 1.4 Hippocampal-Based Generalisation

In a recent perspective article, Zeithamova and Bowman [73] review evidence that the hippocampus contributes both specific memories to generalisation decisions, and also forms generalised representations that integrate information across experiences. They propose four candidate mechanisms, illustrated in Fig 2, that may underpin such “hippocampal-based generalisation”; (a) separate encoding of individual events without integrated representations – generalisation here occurs on demand depending on the task at hand, (b) integrated encoding (e.g. if an appropriate schema, or for us ‘‘cluster”, exists) or separate encoding (if not), (c) initial separate encoding of events that are linked over time due to task demands to form integrated encodings, (d) simultaneous separated and integrated representations.

**Figure 2:**
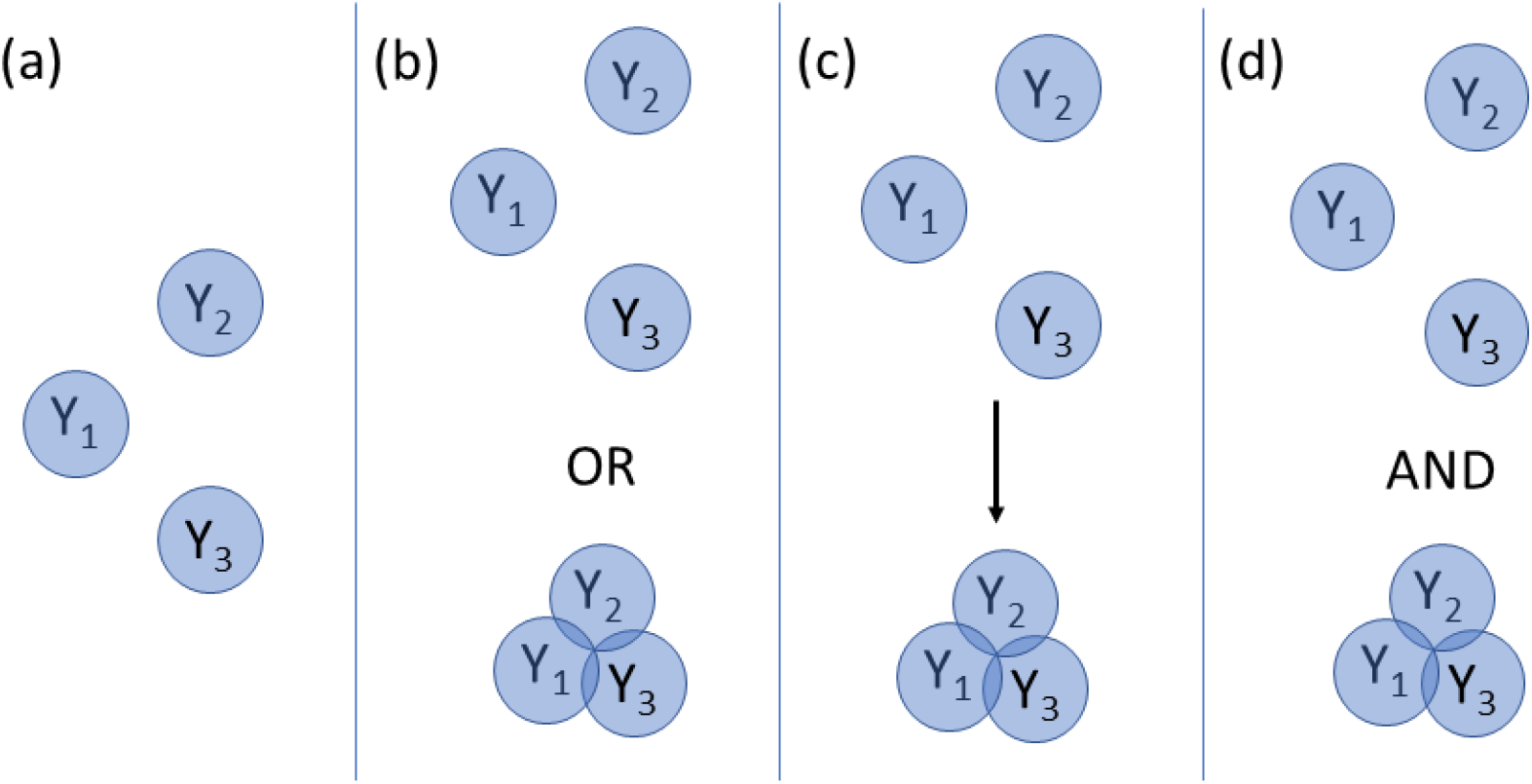
Hippocampal-Based Generalisation. Zeithamova and Bowman propose several different views as to how related experiences (*Y*_1_,*Y*_2_,*Y*_3_) may be represented in memory to support generalisation across experiences: (a) integrated memories are not formed. Alternatively, integrated memories may form (b) instead of separated memories, (c) from initially separated memories, or (d) alongside separated memories. After Fig 5 in [73].

Type (a) computational models include approaches by Bornstein and Daw [17], Viejo et al.[19] and Kumaran et al. [42, 41] (plus see refs in [73]). Although CMM is a joint probability model, the conditional probability of categories given stimuli is computable via pattern completion (see section below). With this in mind, CMM can support decision making using a type (a) mechanism. However, because the clusters within CMM are optimised based on the joint probability rather than the conditional probability, the clusters are not best located, sized, or oriented for categorisation.

The CTM, however, is optimised for decision making. Its clusters are initially set identical to the CMM but then clusters are merged so as to maximise the conditional probability (see section on “cluster merging” below). This merging effectively integrates over multiple related experiences creating larger clusters that are optimised for decision making. This could correspond to a type (c) mechanism if CTM replaces CMM or type (d) if the two representations coexist (although the CMM would be created first). This paper can be viewed as providing a response to the call for more precise definitions of memory schemas [29] and algorithms implementing the various types of hippocampal-based generalisation [73]. We return to the broader literature in the discussion.

### 1.5 Rational Category Models

The Cluster-Based Modelling framework we propose is based on closely related lines of work in probabilistic cognition: Rational Category Models (RCMs) [1] for category learning and Latent Cause Models (LCMs) for reward learning [13, 26]. LCMs and RCMs both provide joint density models over variables of interest and use the same nonparametric cluster-based approach to compute the density using pattern completion (see below). Both derive the conditional densities required for decision making from the joint density. As this paper focusses on category learning we delay discussion of the LCM to the discussion section.

The RCM is a classic model of category learning from the behavioural psychology literature which has been revisited in light of developments in Bayesian Nonparametric Models [57, 58]. It is an adaptive clustering approach in which the number of clusters grows with the number of exemplars presented. By integrating over clusters, it provides a “joint density” over sensory stimuli and category labels. The conditional probability of categories given stimuli that is required for decision making can then be computed by pattern completion (see below).

Our approach inherits a number of attractive properties from RCMs and LCMs. RCMs embody a natural trade-off between the exemplar and prototype models of categorisation that have been proposed in the mathematical psychology literature. RCMs with multiple clusters each based on a single exemplar correspond to exemplar models, whereas RCMs with a single cluster correspond to prototype models (see Figure 2 in [58]). Additionally, RCMs can learn multiple clusters/prototypes for each category and so implement nonlinear decision making. The size and number of clusters is determined by model parameters and adapts to the nature of the underlying categorisation problem.

Following both RCM and LCM the CBMs defined in this paper use an online learning algorithm that requires a single pass through the data. For RCMs and LCMs this has has previously been implemented using inference algorithms based on Maximum a Posterior (MAP) [1], Particle Filtering [58], or Expectation-Maximisation (EM) [27] principles. In this paper, however, we instead use a Variational Inference (VI) approach [4, 11]. This is necessary because cluster consolidation (see Fig 1) is based on a model comparison metric, the model evidence, a quantity readily provided by VI but not the other algorithms.

## 2 Methods

We take a top-down modelling approach as with other probabilistic models of cognition [33]. The first section describes general properties of Cluster Based Models including recognition, learning, inference, pattern separation, pattern completion and generative replay, all illustrated using a simple example with two sensory inputs. The following section then presents a formal specification of our probabilistic model, with subsections describing the priors, likelihood and Variatonal Inference algorithm. This is followed by a section describing the cluster merging procedure and the description of a behavioural task and benchmark data which are used in the results section.

### 2.1 Properties of Cluster Based Models

A Cluster Based Model (CBM) is simply a mixture model, but where each data vector *y* has components arising from multiple sources *y* = [*x*_1_,*x*_2_,…, *x_i_*,…,*x_I_*]. For example, *x*_1_ might be a vector of visual input, *x*_2_ auditory input, *x*_3_ interoceptive input, *x*_4_ category label or reward, and *x*_5_ a task label. Each data vector *y* is assigned to a discrete cluster, *z*, a latent variable in the model. The likelihood of a data vector is assumed to factorise over the components (or groups of components). That is

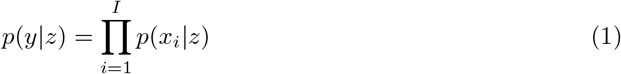

The joint density over an observation is then

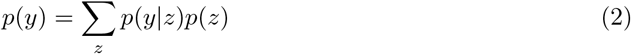

and the probability distribution over latent variables given an observation is

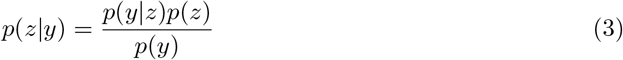

More concretely, in this paper we have *y* = [*u, c, s*] with sensory stimuli *u*, assumed to be continuous and multivariate, category labels *c*, assumed discrete, and task labels *s* also assumed discrete. We could use the term Mixture Model instead of CBM but the most important feature of these models is that they cluster data, hence we prefer the term CBM. Before getting into the details of the model we take a step back to consider some general properties. For the moment, we ignore the category and task labels to focus on a simple Gaussian mixture model of bivariate sensory data, *u*. This data may be thought of as the firing rate of two cells, sampled over different trials or time intervals, impinging on hippocampus. Tutorials on Gaussian mixture models are available in standard textbooks [9, 10]. Figures 3 and 4 illustrate several characteristics of this model which also follow over to the more general CBMs. The data *u* are modelled as arising from the mixture

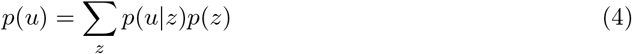

just as in Eq 2 but with *y* = *u*. Each of these distributions depend on model parameters *θ* (the means, covariances and prior probabilities of each cluster) for the two clusters *z* = {1, 2}. As shown in the central panel of Fig 3 this model has been trained on the data points shown as small blue crosses. When a new data point *u* is presented, depicted by the black cross, Eq 4 can be used to compute a recognition score *p*(*u*). The process of “recognition” is illustrated in the left panel of Fig 3 (the similar height of the two bars over *z* reflects the similar number of exemplars belonging to each cluster – the prior probability). Bayes rule can then be used to compute the posterior probabilities of cluster membership

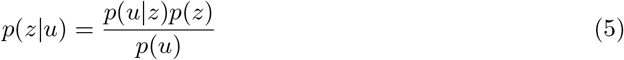

**Figure 3:**
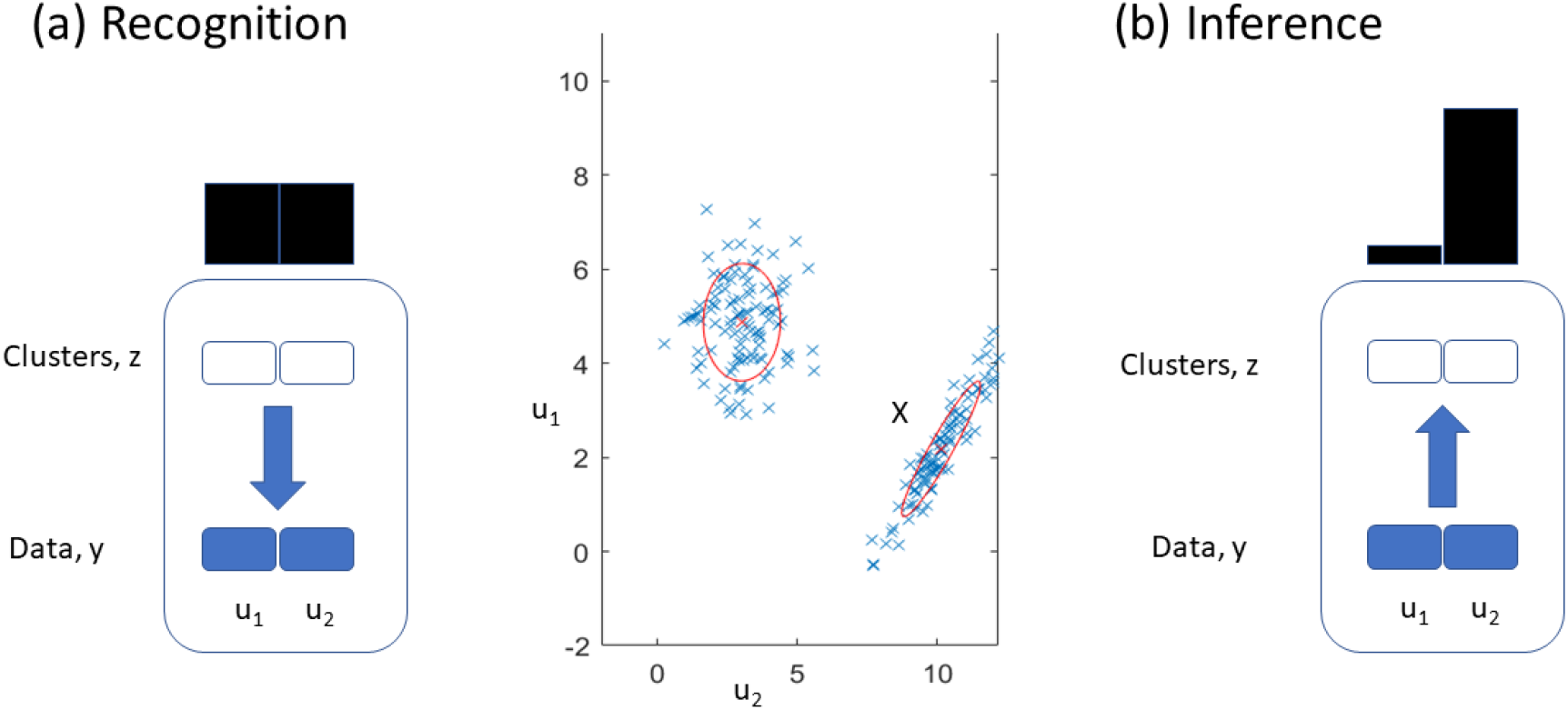
Recognition and Inference. In the recognition step, a new data point *u* (large black cross in central panel) is assigned a recognition probability *p*(*u*), based on the prior cluster probability *p*(*z*) (bars near top of left panel), and the likelihood *p*(*u|z*). In the inference step, the new data point is assigned to a cluster, with the posterior cluster probability bars in the right panel, *p*(*z|u*), indicating that *u* is soft-assigned to the bottom right cluster. Learning occurs when the parameters (e.g. cluster priors, means and covariances) are updated to accommodate the new data point. A new cluster can be created if the new data point is sufficiently novel.

This “inference” step is illustrated in the right panel of Fig 3 with the height of the bars denoting *p*(*z|u*) for *z* = {1, 2}, here indicating that the new data point most likely belongs to the bottom-right cluster. The parameters of this cluster can then be updated to accommodate this new point. In a non-parametric approach, if the new data point is not sufficiently close to either cluster then a new cluster is created. Parameter updates and (potential) cluster creation constitute the “learning” step.

The left panel of Fig 4 illustrates “cued recall” or “pattern completion”. Here a partial cue is presented to the model, *u*_1_ =6. Eqs 4 and 5 are used to compute the cluster belonging probabilities, but adapted because only *u*_1_ is observed

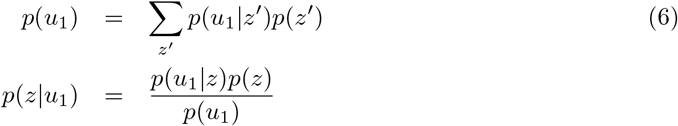

here indicating that the top-left cluster is most likely. Pattern completion is then based on the predictive density

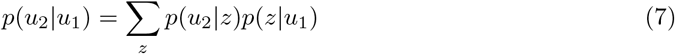

**Figure 4:**
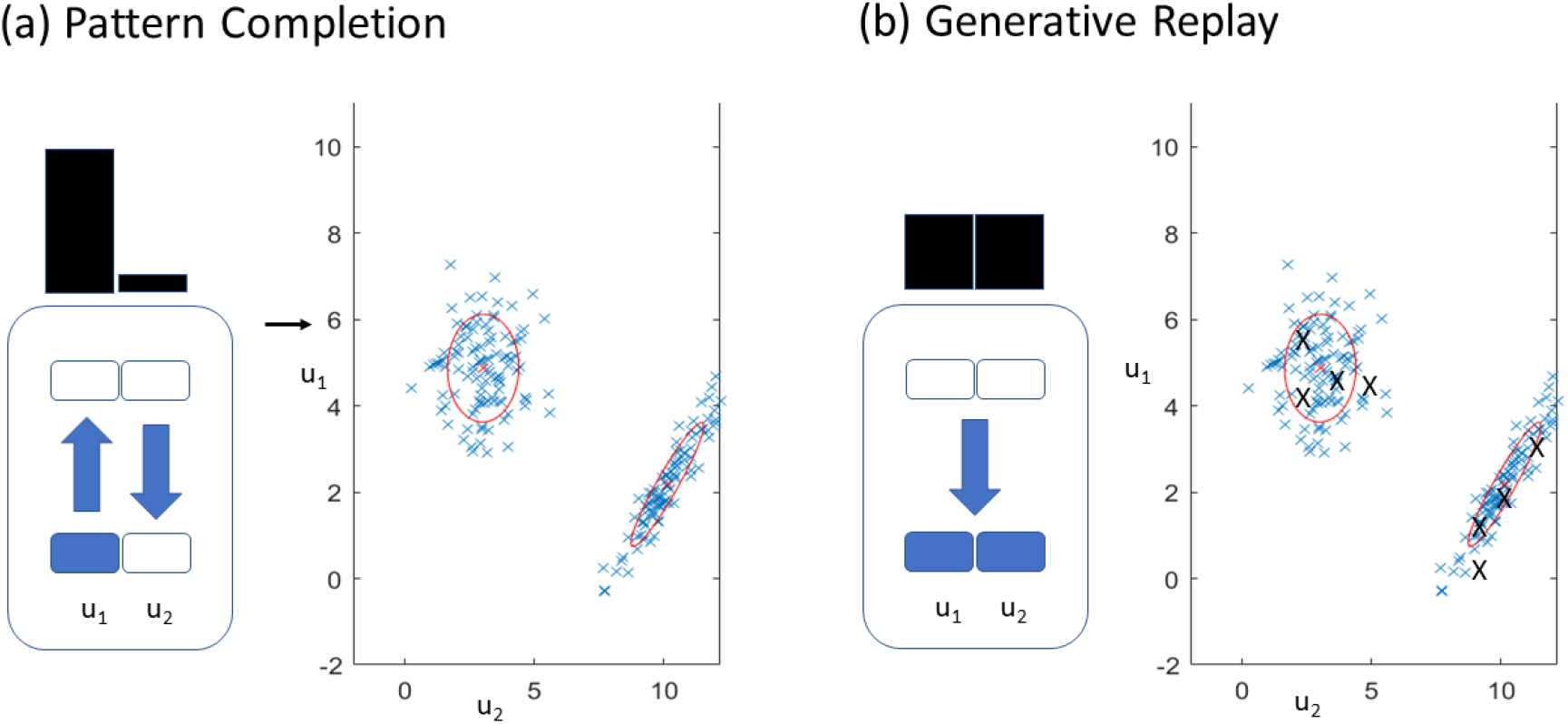
Pattern Completion and Generative Replay. In panel (a) a partial cue is presented, indicated by the arrow at *u*_1_ = 6, and the posterior distribution over clusters, *p*(*z|u*_1_), indicates the top-left cluster to be the likely cause. Pattern completion is implemented by computing and then sampling from the density *p*(*u*_2_|*u*_1_) as shown in Eq 7. Panel (b) demonstrates generative replay in which new data points, shown as large black crosses, are sampled from the joint density, *p*(*u*). These new data have the same statistical structure as the original data.

The right panel of Fig 4 illustrates “free recall” or ‘generative replay”. Here, we are simply drawing samples from the joint density *p*(*u*). This takes place in a two step approach, which correspond to the two terms in Eq 4; the first samples latent variables from *p*(*z*), the second samples observations from *p*(*u|z*). Loosely, this may correspond to the production of ripples in CA3 and the resulting CA1/entorhinal activity. This produces samples shown as black crosses in Fig 4 which have the same statistical structure as the original data. We note that the above properties and related computations also apply to any generic “latent variable model”, that is, a model with a single layer of hidden variables, and a layer of observed variables. If the latent variables are continuous then the sums in the above equations are replaced by integrals.

Further characteristics that are relevant for CBM as a model of hippocampus are that (i) cluster-based representations are localised in stimulus space (c.f. place cells), (ii) the distribution over latent variables is sparse (the posterior over clusters *p*(*z|y*) – see Eq. 3 – will only have significant mass for a small number of clusters), (iii) because the data likelihood factorises over the source likelihoods (Eq 1) then cluster activations, *p*(*z|y*), are computed using a conjunction (followed by normalisation) and may reflect conjunctive representations in hippocampus [16], (iv) clustering is a natural mechanism for segmenting experience into similar chunks [49].

Another characteristic of CBM that we wish to emphasise is that category labels (and in a companion paper, rewards) are treated in the same way as sensory inputs – they’re just another variable over which to compute the joint density. Having both rewards/categories and sensory inputs as part of the same data vector is also inherent to the Gluck-Myers hippocampal representation theory [30]. This has been noted to usefully bias latent representations (in our case clusters) to also reflect rewards (or categories) and has been adopted as part of an unsupervised predictive model of memory in a recent machine learning architecture [68]. It is also a key component of Rational Category (see above) and Latent Cause Models (see discussion). Having a generative model that includes categories allows generative replays that are specific to each category, a feature that has been shown useful for continual learning applications in AI [67].

Looking at Figs 3 and 4 it may appear that all detail about individual items, the small blue crosses in the central panels, is lost. This isn’t the case, however, as we are viewing *u*_1_ and *u*_2_ as spike rates generated in response to sensory input. It makes no sense to have a hippocampal precision that is higher than the precision of incoming signals. Generally, CMM will use an encoding precision which is high enough for episodic memory and generative replay, whereas CTM recursively merges clusters to produce a model that is optimised for categorisation (see Fig 9 below).

There are characteristics of episodic memory and hippocampal architecture, however, that are distinctly lacking in CBM. First, if such a model were applied to sensory data arriving directly from primary sensory cortices it would not be able to handle the *high dimensionality.* But, of course, episodic memory systems and hippocampus do not receive primary sensory input either. We envisage that the CBM architecture could play a useful role if its input were from the sort of compact, multimodal representations that are available in association cortex. We return to the issue of dimensionality in the discussion. Additionally, when using CBM to classify handwritten digits from the MNIST database, we use a dimensionality reduction algorithm of the sort though to be supported by neocortex (see [34] for a similar approach in the context of working memory).

Second, a fundamental characteristic of episodes is their *sequential nature* and one of the primary characteristics of memory recall is its time- and order-dependency. CBMs as described in this paper are not temporal models and so cannot capture these order effects. However, recent work in the related Latent Causal Model framework proposes a model of memory modification [27] by changing the prior *p*(*z*) to reflect only recent cluster assignments (rather than from all assignments to date). This can implement power-law forgetting functions of the sort found in empirical studies. We return to the broader issue of temporal structure in the discussion.

### 2.2 Cluster-Based Model

This section describes the mathematics underlying the cluster-based model. This is a joint density model of sensory inputs, category labels and task labels. The model can learn from data presented sequentially, e.g. one trial at a time. The algorithm is derived in such a way that it also permits batch learning (for applications beyond the scope of this paper). It uses a variational inference approach where data is assigned to clusters in a variational E-step and model parameters are updated in a variational M-step. It allows for semi-supervised learning where category labels are provided for only a subset of trials. This section also describes the cluster consolidation process which is based on moment-matching and a variational approximation to the evidence of the conditional density model.

Broadly, we consider a sequential learning or “data streaming” setting in which *b* = 1..*B* blocks of data each contain *T* data points. For most of the applications in this paper we have *T* =1 data point, or experimental trial, per block and the E- and M-Steps described below are iterated for each single data point.

We are given a set of categories *C* = [*c*_1_, *c*_2_,…,*c_T_*], task variables *S* = [*s*_1_,*s*_2_,…,*s_T_*], and d-dimensional input cues *U* = [*u*_1_,*u*_2_,…,*u_T_*], and a (within data block) cluster-based generative model with latent variables *Z* = [*z*_1_, *z*_2_,…, *z_T_*] as shown in Fig 5. The latent variables *z_t_* indicate the cluster. The joint probability of data and latent variables is

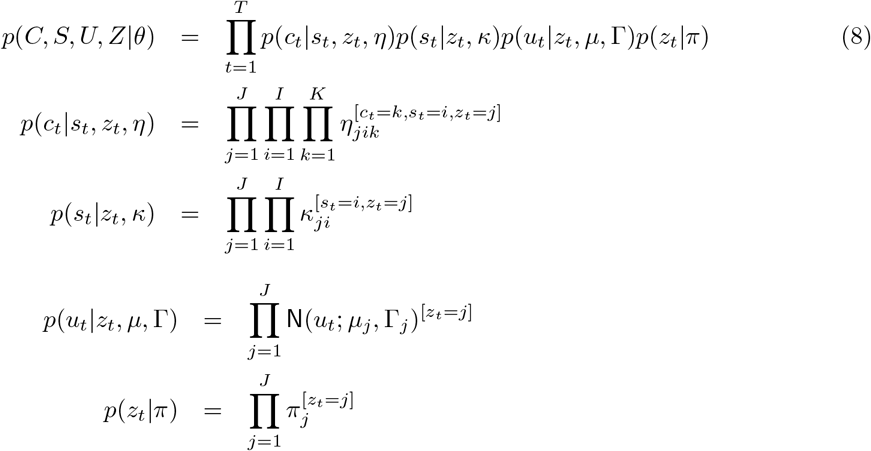

where the notation [*a* = *b*] returns 1 if the condition *a* = *b* is satisfied, and 0 otherwise. This is known as an Iverson bracket. Data is collectively written as *Y* = {*C,S,U*} and model parameters as *θ* = {*η,κ,μ*, Γ, *π*}. Here, N(*x; m,* Λ) is a multivariate normal distribution with mean m and precision matrix Λ, and in what follows Ga(*x; b, c*) is a Gamma density with shape *c* and inverse scale *b*, Dir(*π*; λ) is a Dirichlet density with count parameter λ, and W(Γ; *a, B*) is a Wishart density over matrix Γ. These distributions are defined in a separate document [53].

**Figure 5:**
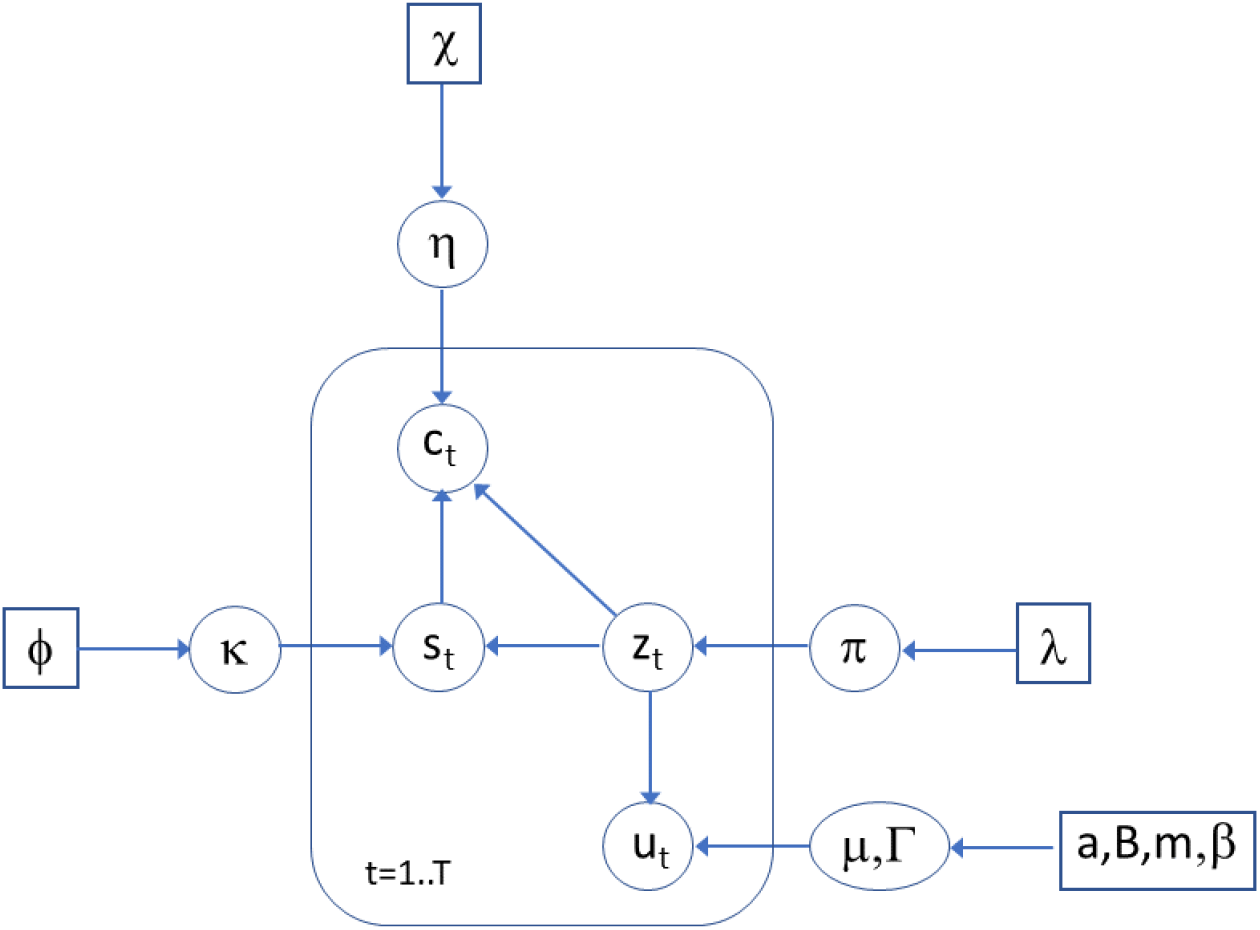
Generative Model. We are given a set of categories *C* = [*c*_1_,*c*_2_,…,*c_T_*], task variables *S* = [*s*_1_, *s*_2_,…, *s_T_*], and sensory input cues *U* = [*u*_1_, *u*_2_,…, *u_T_*], and a cluster-based generative model with latent variables *Z* = [*z*_1_, *z*_2_,…, *z_T_*] where *z_t_* indicates the cluster that is active on trial *t*.

#### 2.2.1 Priors

The prior on model parameters is

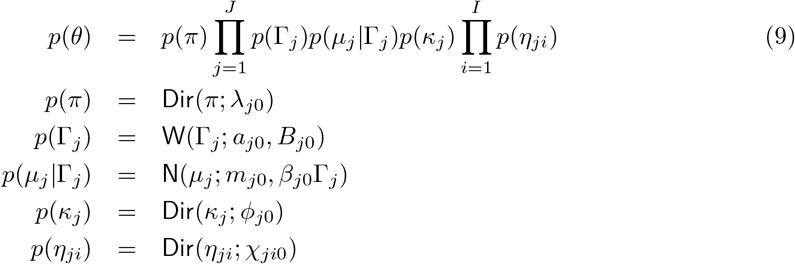

Together, the prior *p*(*μ_j_*, Γ_*j*_) is a Normal-Wishart density. The mean prior covariance is 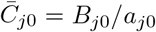 and is an important quantity as it sets the precision of stimulus encoding. As we shall see in the Results section below, a high precision is needed for online learning in the CMM.

#### 2.2.2 Approximate Posterior

Generically, a variational approximation to the posterior can be derived by assuming the factorization

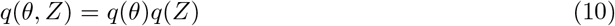

The parameter posterior *q*(*θ*) and latent variable posterior *q*(*Z*) can then be obtained by maximizing the variational objective function [2, 4]. This is also known as the ‘negative variational free energy”, *F*, which provides an approximation to the model evidence. The generic expression for F and procedures for obtaining *q*(*θ*) and *q*(*Z*) are outlined in Appendix A and B. For the parameters this gives

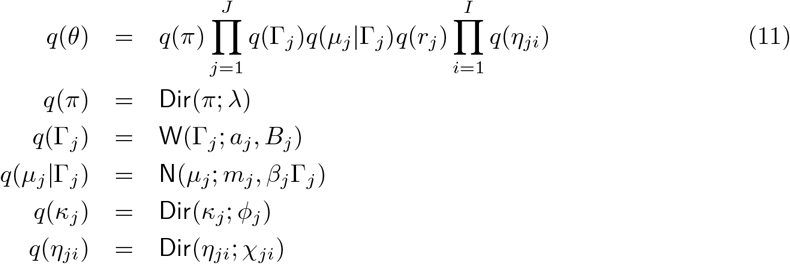

Because the approximate posteriors have the same functional forms as the priors (‘conjugacy’), the algorithm can be applied to blocks of sequentially arriving data (see ‘Sequential Inference’ below). For the latent variables we have

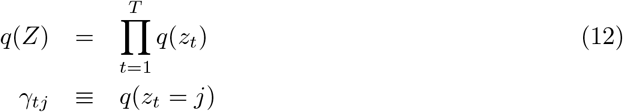

where *γ_tj_* is the probability that component *j* is responsible for category, task and cue data at time *t*. The approximate posteriors *q*(*Z*) and *q*(*θ*) are then updated in what are referred to as the variational E- and M-Steps. The E- and M-Steps are described in Appendix B and are iterated until *F* reaches a maximum.

#### 2.2.3 Sequential Learning

The algorithm we have described can be applied to data arriving sequentially in blocks with Y denoting the data arriving in each block. Before the first block the priors can be set as follows

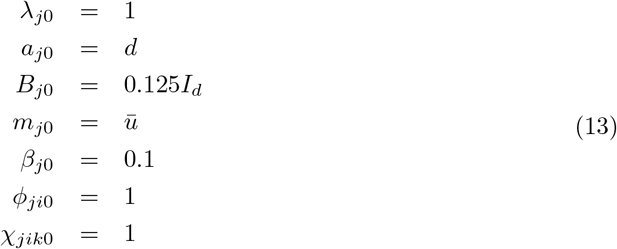

where 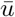 is the mean cue vector. Strictly, this is not a quantity that can be known in advance but in a neural processing context we envisage that firing rates are normalised within a certain known range. In subsequent blocks the prior parameters are set to the posteriors from the previous block. That is,

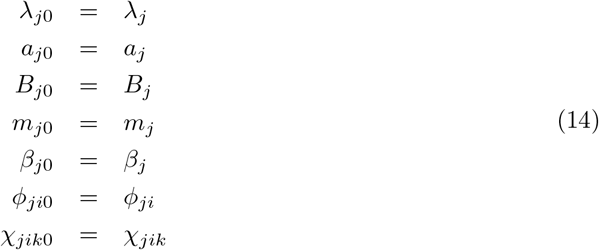

For the behavioural learning simulation we use Sequential Inference with a block size of a single sample. The same algorithm can be used with a larger block size which may accommodate, for example, a more generous working memory capacity. For machine learning applications the block size could increase to the size of the whole data set, though could be reduced in data streaming or online learning contexts.

#### 2.2.4 Cluster Creation

Central to the definition of Rational Category Models (RCMs) is a nonparametric approach in which the number of clusters can increase in proportion to the number of data points. We implement this in the context of sequential learning. The following approach allows the number of clusters to increase by one at each sequential learning step, and is perhaps most useful for when the new batch of data contains a single example (as with RCM).

At each sequential learning step we fit two models, one with the original number of clusters, *J*, and an augmented model with *J* +1 clusters. For the latter model, the new cluster is initialised by setting parameter values equal to prior values (see Eq. 13) except for the stimulus mean, *m_j_*, which is initialised to the new cue value (or mean thereof if the new batch of data contains multiple samples). The prior over clusters in the original model is as defined above, that is

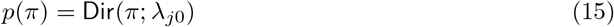

For the augmented model we use the same *λ*_*j*0_ parameters for *j* = 1..*J* and for the new state

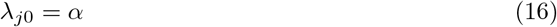

where *α* is a parameter that determines the probability with which new clusters are created. For the simulations in this paper we use *α* = 10. Finally, after model fitting, if the posterior probability of cluster *J* + 1 is greater than for any other cluster we accept the new cluster, otherwise it is rejected and the original model is returned. This Maximum A Posteriori (MAP) criterion was also used in the original RCM [1].

#### 2.2.5 Semi-Supervised Learning

The model we have proposed can be used for semi-supervised learning, that is, for data sets in which category labels and/or task labels are not provided on every trial. On trials for which this information is absent, the task and/or category terms in Eq. 23 (the first two terms) will have a zero in the exponent for all j, resulting in the corresponding terms being constant. The cluster posteriors will then solely reflect contributions from other terms (i.e. from the remaining data types provided on that trial). If category and task labels are absent from all trials then the model reduces to a Gaussian Mixture Model (over cues).

### 2.3 Cluster Merging

We define a procedure for merging clusters based on previous proposals in the literature. We need (i) a *merging procedure* for computing the parameters of the merged from the unmerged model, (ii) an *evaluation criterion* for deciding whether or not to proceed with the merge, and (iii) a *proposal heuristic* for proposing which clusters to merge. We refer to the merging process as “cluster consolidation”. Here, we take consolidation to mean the process of “combining a number of things into a single more effective whole” as opposed to ‘making something stronger” which is rather reflected in the various synaptic consolidation processes involving tagging and protein synthesis [56].

#### 2.3.1 Merging Procedure

Based on previous work by Richardson and Green [55], Zhang et al. [74] proposed a moment-matching principle for merging clusters in multivariate Gaussian Mixture Models whereby the zeroth moments (mixing coefficients), first moments (means) and second moments (covariances) of the merged and unmerged clusters are equated. Here, we also extend this principle to the multinomial distributions over tasks and categories. Using this approach (and equating degrees of freedom for *α_j_, β_j_, λ_j_*) the parameters of a merged cluster, *n*, created by combining clusters *j* and *j*’ are given by

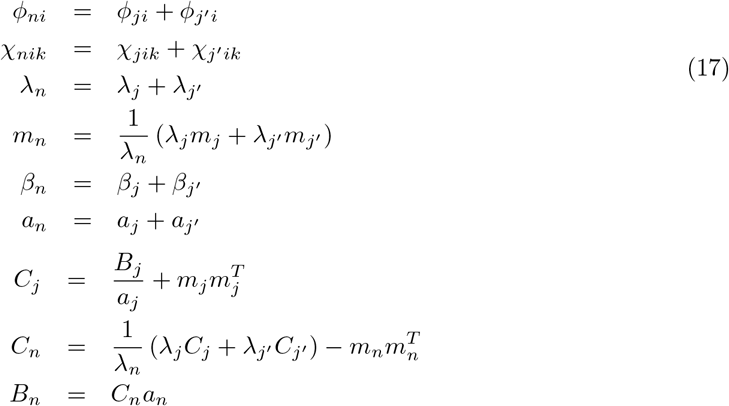

#### 2.3.2 Evaluation Criterion

Following Beal [4] we will use the variational approximation to the model evidence, *F*, as our evaluation criterion. However, as explained in the introduction this will be the evidence of the conditional density model, *F^C^*, rather than the joint density model, *F^J^*. As derived in the appendix, we have

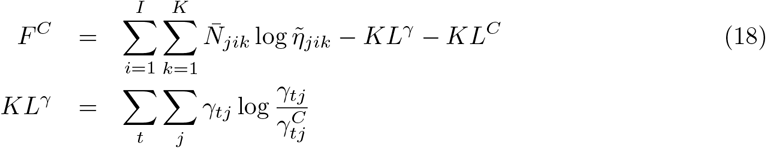

where the first term in *F^C^* reports correct classification likelihood, *KL^γ^* penalises CTMs in proportion to how different cluster assignments are to their corresponding CMMs, and *KL^C^* penalises models with more clusters. As with most model comparison criteria, *F^C^*, is comprised of an accuracy term (first term above) and a complexity term (the two KL divergences). The quantity *F^C^* is computed using *N_s_* data points sampled from the CMM. For the empirical work in this paper we set *N_s_* to be the number of samples used to create the CMM. The E-Step equation is used to compute 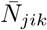 and *γ_tj_* on this data, and we use Eq 38 to compute 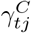. This then provides a sample-based approximation to *F^C^*. Effectively, we are using generative replay to compute the model evidence. Informally, if clusters can be merged this will reduce model complexity, and if this doesn’t adversely affect model accuracy then the merge step will be accepted.

We propose that clusters *j* and *j*’ are merged to form cluster *n* where the parameters of cluster *n* are set as described in the previous section. A subtlety in the context of Sequential Inference is that, to compute *F^C^*, we use the prior before any data have been encountered e.g. the values in equation 13, rather than the prior before observing data in the last block. This is because we wish model comparison to be based on the whole data set, rather than just the data in the last block. If 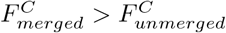 then the proposal is accepted.

#### 2.3.3 Proposal Heuristic

Because most pairs of clusters will be poor candidates for merging we employ a heuristic for proposing which pairs {*j, j*’} to merge. Clusters are first designated based on their maximum posterior category. Euclidean distances are then computed between within-category cluster means and ranked accordingly. Rankings are then interleaved across clusters so that an ordered cluster pairing is produced e.g. closest category-1 pair, closest category-2 pair, closest category-3 pair, next-closest category-1 pair, next-closest category-2 pair etc.

#### 2.3.4 Recursive Merging

The merging approach is applied recursively such that, after a successful merge, the resulting model is fed back into the merging subroutine (see above). If an unsuccessful merge is encountered we proceed by trying to merge the next highest ranked cluster pair. This continues up to a maximum of *N_cand_* = 2*K* proposed cluster pairs where K is the number of categories.

### 2.4 Behavioural Data

In the results section we fit CBMs to task data from a recent behavioural experiment (chapter 4 of [47]) in which participants were required to learn the probabilistic mapping shown in the right panel of Figure 6. This was an internet-based study with N = 100 participants recruited from the University of East Anglia. In each trial participants were presented with a fixation cross, a cue appearing on screen, and were asked to predict whether the cue predicted “Sun” or “Rain”. The experiment comprised a 120-trial training epoch, a 120-trial testing epoch (Test1), a 5-minute period of Quiet Wakefulness and a second 120-trial testing epoch (Test2). During training, feedback was provided indicating if the participants response was correct or incorrect. Importantly, a distinct set of cues were presented during testing (see left panel Figure 6) for which no feedback was provided. Overall, the test set performance (a measure of generalisation ability) improved from Test1 to Test2. This improvement was significantly nonzero, and significantly greater than a control group who performed a distractor task in place of the period of Quiet Wakefulness (see Figure 6 in chapter 4 of [47]). The results section shows how cluster consolidation is consistent with these improvements in generalisation.

**Figure 6:**
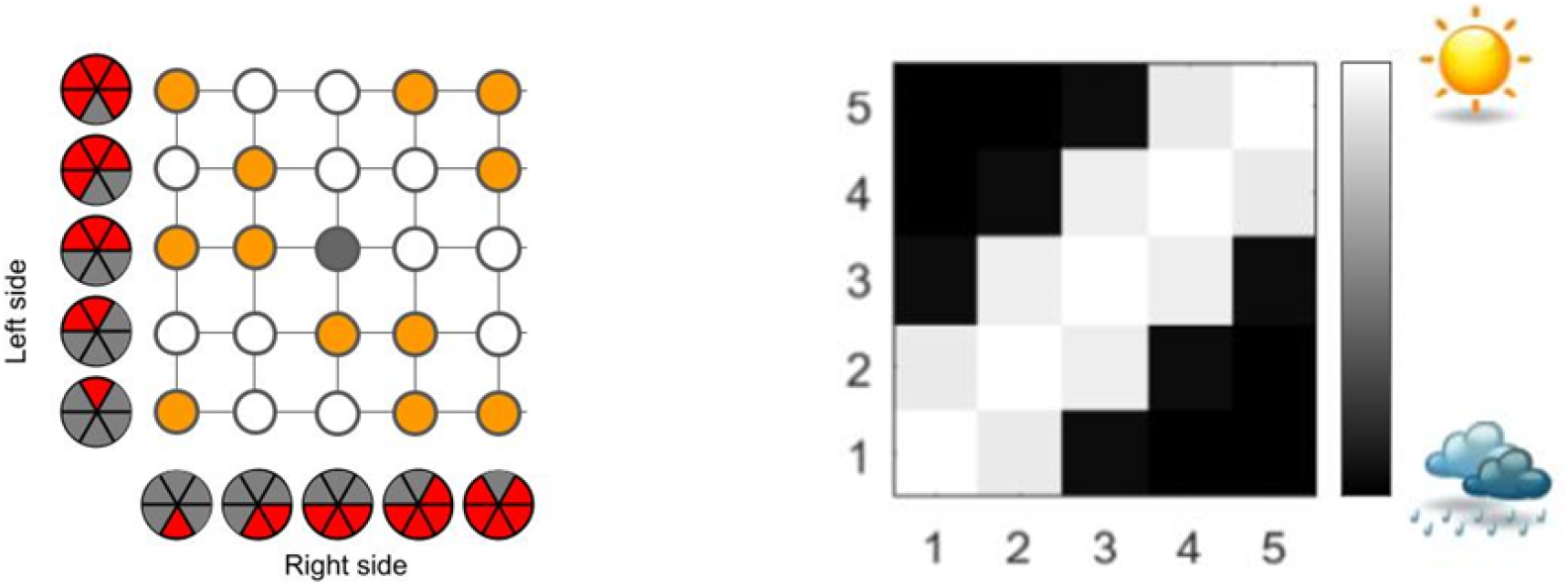
Behavioural Experiment. Left Panel: Yellow indicates cue combinations presented during training and testing, and white indicates those presented during test only. The single grey combination was not presented. **Right Panel:** The gray-scale corresponds to the probability of reward when deciding ‘Sun’. The numbering 1 to 5 on x- and y-axes indicates the number of slices in the right and left pies respectively. Thus, if the number of slices is similar, participants should choose ‘Sun’.

### 2.5 Benchmark Data

A useful constraint on a model of human memory-based cognition is that it should provide good performance of complex categorisation tasks. In the results section we therefore fit CBMs to a number of benchmark classification tasks from the Machine Learning literature, as listed in Table 1. Some of these data have unequal numbers of exemplars in different categories as shown in the table. Three of these data sets are from the UCI Repository https://archive.ics.uci.edu/ml/datasets.php and were chosen because the input stimuli are continuously valued (the cluster-based model, as presently formulated, requires u to be continuous rather than discrete).

**Table 1:** Benchmark Data. The columns indicate the number of data points *T*, dimensionality of input stimuli d, number of categories K, and the proportion of exemplars in of each category.

| Data Set | T | d | K | Proportions |
| --- | --- | --- | --- | --- |
| Iris [20] | 150 | 4 | 3 | Equal |
| Ecoli [36] | 336 | 7 | 4 | [0.47,0.25,0.11,0.17] |
| Ionosphere [61] | 351 | 34 | 2 | [0.36,0.64] |
| MNIST [43] | 1000 | 20 | 10 | Equal |

The remaining data set is the MNIST character recognition data comprising 28 × 28 pixel images of the characters zero to nine [43]. These images were compressed into a 20-dimensional code using a Variational Autoencoder [40]. Although the original images are binary, the elements of the compressed code are continuously valued. The encoder comprised two blocks of 2-D convolution and restricted linear (“ReLU”) layers, and similar for decoding, and was trained on all 60000 training images using the ADAM optimizer as described in the Matlab Deep Learning tutorial https://uk.mathworks.com/help/deeplearning/. In our simulations below we used the first 1000 images (100 of each character).

## 3 Results

In what follows the first two subsections report results on the behavioural data, and the second two on the benchmark data. Unless otherwise stated, the results in the sections below were obtained using the following model parameters: *β*_0_ = 0.1, *a*_0_ = 2, *λ*_0_ = 1, *α* = 10. For all of the simulations we use sequential inference with a block size of a single sample. We define the encoding scale, *σ_enc_* as the prior mean standard deviation in each stimulus dimension i.e. the expected width of the Gaussians in the CMM. Given that the mean prior covariance is *C*_0_ = *B*_0_/*a*_0_ this can be achieved by setting 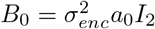.

### 3.1 Online Learning on Behavioural Data

This section examines the dependency of online learning on the encoding scale, *σ_enc_*, and the prior probability of creating a new cluster, *α*. We trained a CMM on 120 training trials of data from the behavioural learning task (see Fig 6). This was repeated for 20 simulations, each with a randomly permuted ordering of cues. These results were obtained with *α* = 10. In what follows, classifications were made by the CMM using pattern completion whereby a category label for a novel cue was inferred using equation 43. The provided category was then used to update the model, and this interleaved process of classification and learning was repeated over all trials.

The average classification accuracy and final number of clusters are plotted in Figure 7. These results indicate threshold effects above which, only a small number of clusters are created and poor task performance results. Conversely, we found only a weak dependence on *α*. Changing *α* to 1 or 100 had only a minor impact on the curves in Fig 7 (with e.g. classification accuracy and number of clusters both increasing a small amount at high *σ_enc_* for *α* = 100).

**Figure 7:**
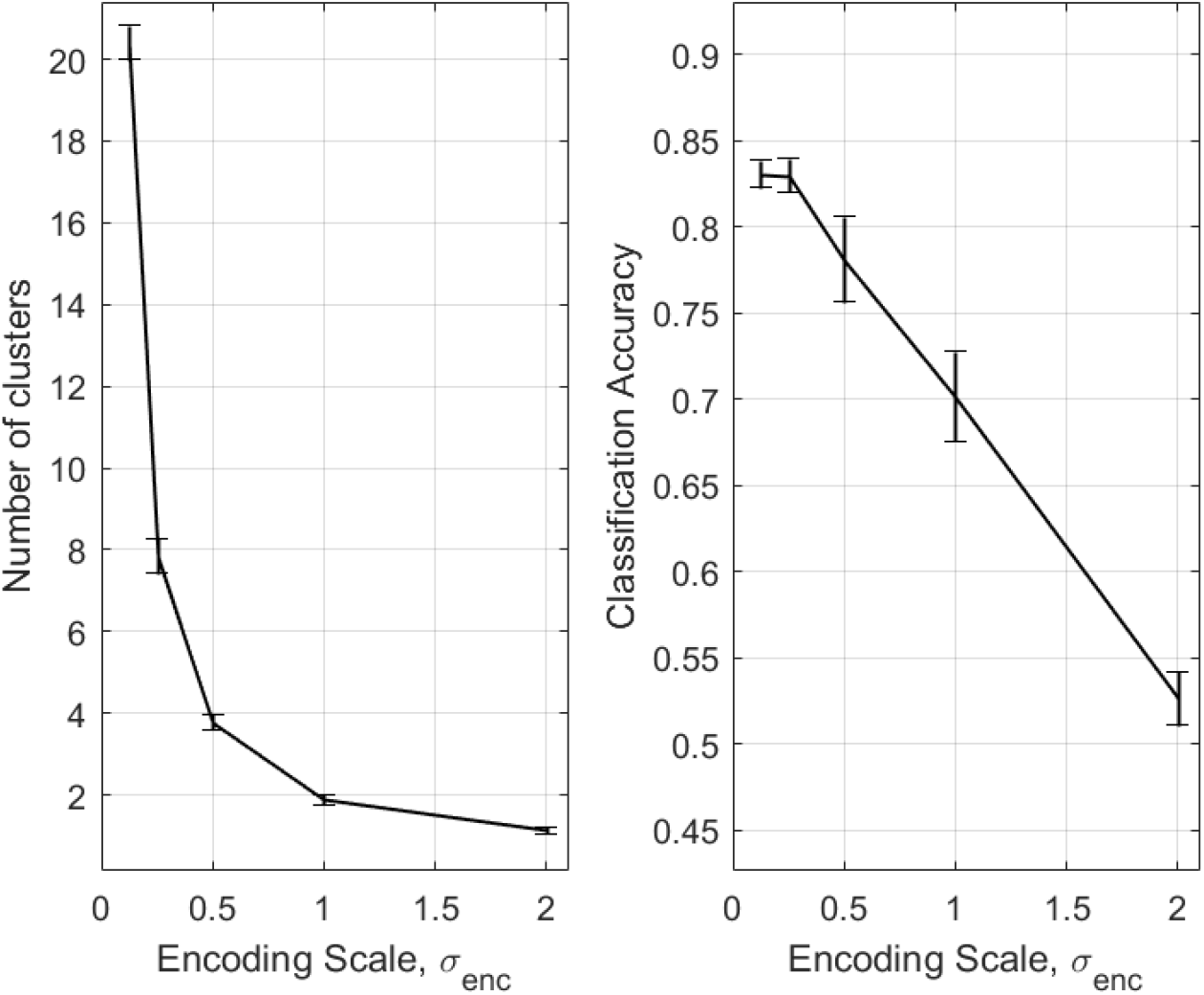
Effect of Encoding Scale on Online Learning. The plots show mean number of clusters and classification accuracy as a function of encoding scale. Error bars denote the standard error of the mean.

### 3.2 Cluster Merging on Behavioural Data

We first trained a cluster based model on 120 training trials of data from the behavioural task comprising 10 repetitions of each of the unique training cues (see Fig 6). We set the encoding scale to *σ_enc_* = 0.25. This was repeated for 120 simulations, each with a randomly permuted ordering of cues. For each simulation we then created two sets of models (1) a set for which we ran cluster consolidation (see section 2.3), and (2) a set for which we did not. We then tested each of the resulting models on 120 test trials with classifications made using pattern completion. To assess memorisation performance we used the training cues, and to assess generalisation we used the testing cues. These two sets of cues do not overlap, as shown in Fig 6. The training cues were learnt with an accuracy of 86% and CMM produced models with an average of 7.36 clusters. Cluster merging reduced this to an average of 2.78 clusters. Merged and unmerged clusterings for a representative simulation run are shown in Figure 8. We can see that the cluster labelling reflects the structure of the behavioural task (see Fig 6). In contrast to the CMM, the CTM representations do not have good pattern separation but provide generalisation across learning episodes. Figure 9 shows a typical sequence of cluster merges.

**Figure 8:**
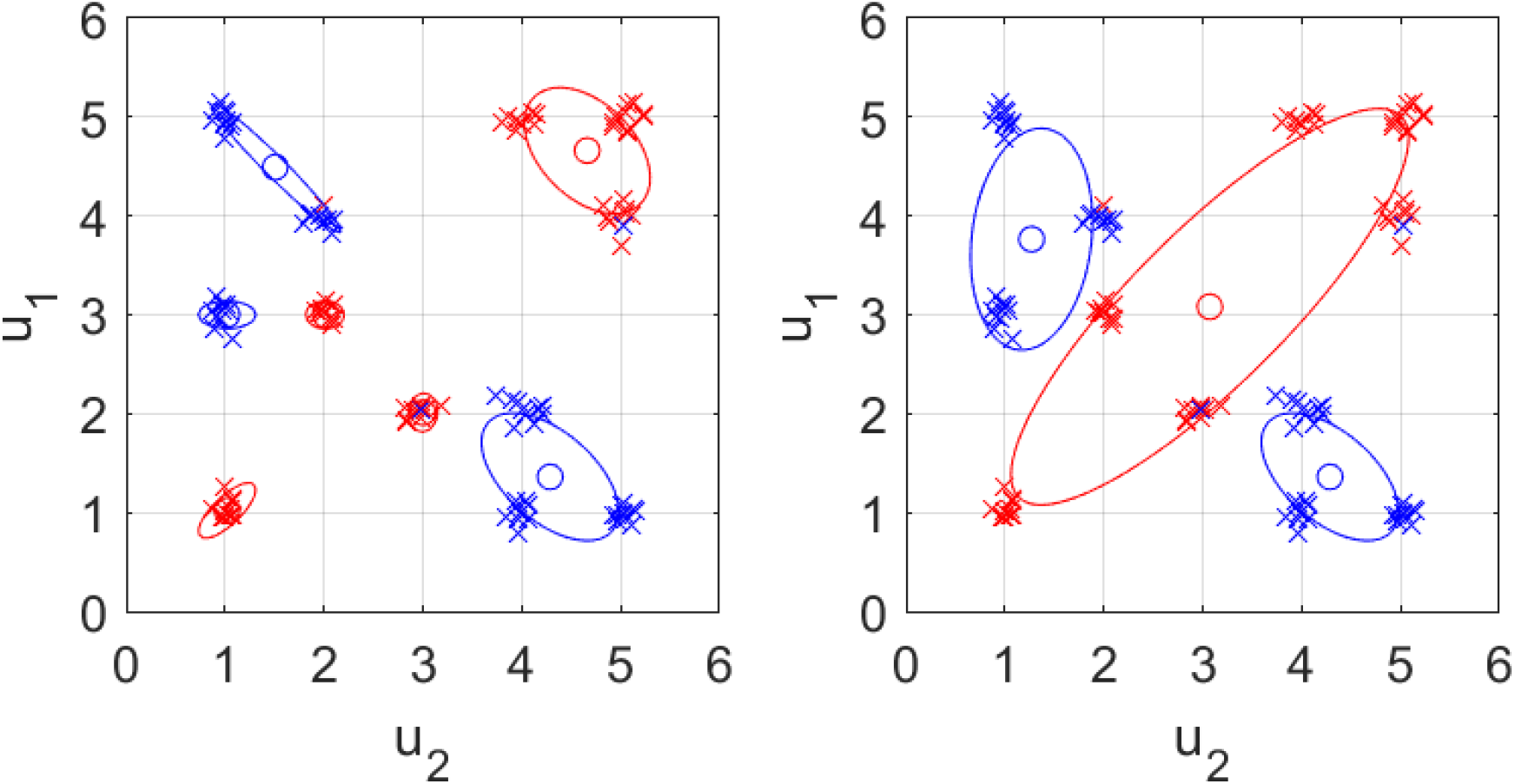
CMM versus CTM Clusters. Clusters produced by CMM (left panel) and subsequent cluster merging (right panel) are shown in red for category 1 clusters (with category probabilities *η*_*j*11_ > *η*_*j*12_) and blue for category 2. The left panel shows the representations within CMM. We can see that unique cues are frequently associated with unique clusters, thus producing strong *pattern separation.* The right panel shows the representations within CTM which have been created through cluster consolidation and exhibit *generalisation across learning episodes.* The data points are shown as crosses (red for category 1) and have been randomly jittered to aid visualisation.

**Figure 9:**
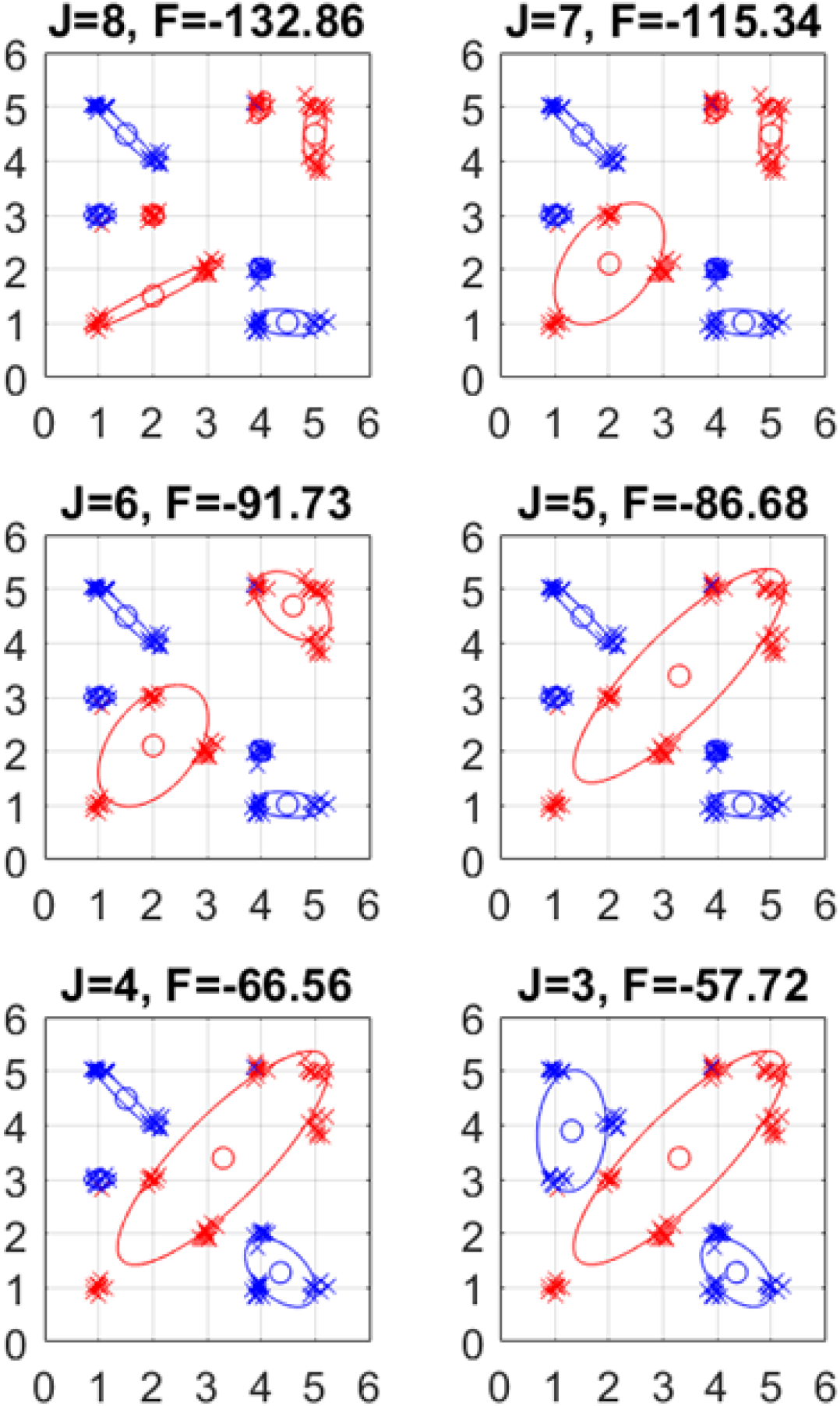
Cluster Merging Process. During online learning, clusters were set to have a reasonably high encoding precision (with *σ_enc_* set to 0.25) resulting in *J* = 8 clusters (top left). Many unique cues have their own cluster, resulting in good pattern separation. This is the representation in the Cluster-based Memory Module (CMM) and provides the starting point for cluster merging. Clusters are merged so as to maximise the evidence of the conditional density model (Eq 40). At each merge step we can see that the (approximation to the) log evidence of this model, *F*, increases. The recursive merging process produces the Cluster-based Task Module (CTM) which has *J* = 3 clusters (bottom right) and has identified the latent structure of the task (see Fig 6 for comparison). The simulation reported in this figure used a different random ordering of cues than for the results shown in Fig 8. The data points are shown as crosses (red for category 1) and have been randomly jittered to aid visualisation.

During the testing phase, accuracy on new cues was 67.9% for merged models, and 53.6% for unmerged. The difference of 14.3 is statistically significant (*t*(119) = 17.9, *p* < 10^-3^). The accuracy on old items was 83.4% for merged models, and 88.8% for unmerged. The difference of 5.4 is statistically significant (*t*(119) = 18.9, *p* < 10^-3^). These results are plotted in Figure 10. The overall accuracy (on new and old cues) was 75.6% for merged models, 71.2% for unmerged. The difference of 4.4 is statistically significant (*t*(119) = 9.83, *p* < 10^-3^). The reduction in classification performance on old items suggests that there is a component of forgetting which is functional; forgetting is necessary to improve future performance.

**Figure 10:**
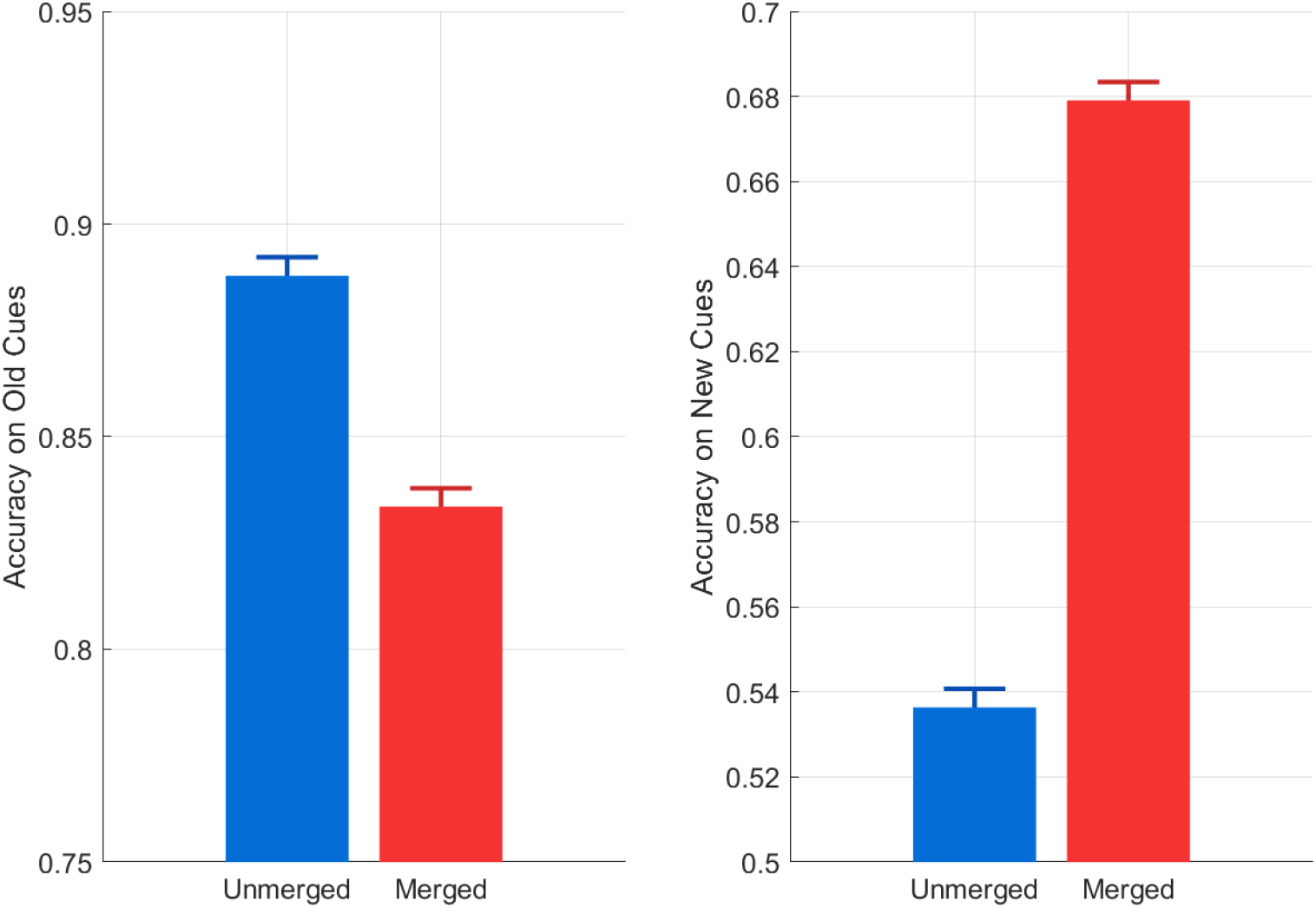
Effect of Cluster Merging on Task Accuracy. Cluster merging improves generalisation (accuracy on new cues), at a cost of memorisation (accuracy on old cues), but overall yields improved performance (see main text).

We also plot the recognition scores in Fig 11 for new and old items from both merged and unmerged models (CMM and CTM respectively). The scores are computed according to Eq 41 in the appendix. The scores plotted in Fig 11 are averages over all stimulus items in the test set. In summary, we find that cluster merging improves generalisation, at a cost of memorisation, but overall yields improved performance.

**Figure 11:**
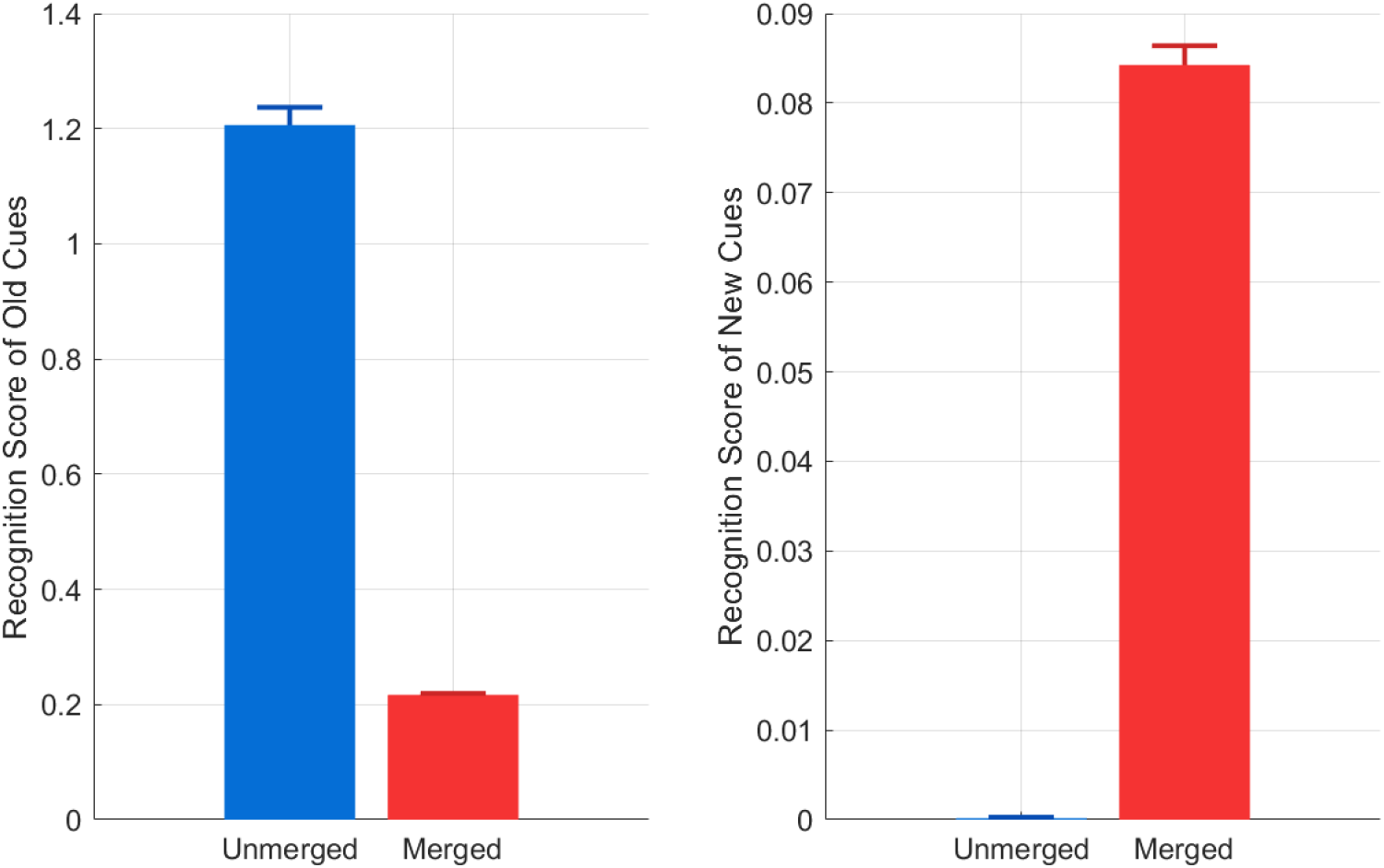
Effect of Cluster Merging on Recognition Scores. The cluster-based model produces recognition scores for each stimulus which are computed as shown in Eq 41. The left and right panels plot these scores for old and new cues, respectively. Blue bars indicate scores for the unmerged model, corresponding to the Cluster-Based Memory Module (CMM), and red scores for the merged model, corresponding to the Cluster-Based Task Module (CTM). CMM has high recognition scores for old items only, whereas CTM has higher recognition scores for new items at a cost of lower scores for old items.

### 3.3 Semi-Supervised Learning on Benchmark Data

The majority of experiences from which humans learn are thought to be unlabelled. In a classification context this means that the majority of stimuli are not paired with category labels. As described in section 12 this is readily accommodated in a CMM. Here, we demonstrate that CMM can be superior to a standard supervised learning approach (which cannot make use of unlabelled data), Softmax Regression [48] (the multiple category generalisation of Logistic Regression). We use the Machine Learning benchmark data. On each simulation run, the data sets were randomly permuted and split into equal sized training and testing sets. Category labels were then removed from a proportion of exemplars. CMM was trained online (to maximise the joint density of stimuli and labels) using a single run through the data set. Softmax Regression was trained only on that proportion of the training data that was labelled (it has no way to make use of unlabelled data). Softmax Regression was trained offline, allowing multiple runs through the data set so as to maximise the conditional density of labels given stimuli. We also report results for online version of Softmax Regression, which allows only a single run through the data. The accuracy on the test set was assessed using the weighted balanced accuracy [32]. This criterion is suitable when categories contain unbalanced numbers of exemplars. It is computed as the sum of category sensitivities weighted by the category frequencies. We ran a total of 40 simulations for each data set and report results in Fig 12. CMM provides better performance for the Iris and Ecoli data but worse for Ionosphere and MNIST. For Iris and Ecoli, CMM provides good performance with even a small proportion of labelled data e.g. 10%. The failure of CMM on the Ionosphere and MNIST data will be addressed in the discussion.

**Figure 12:**
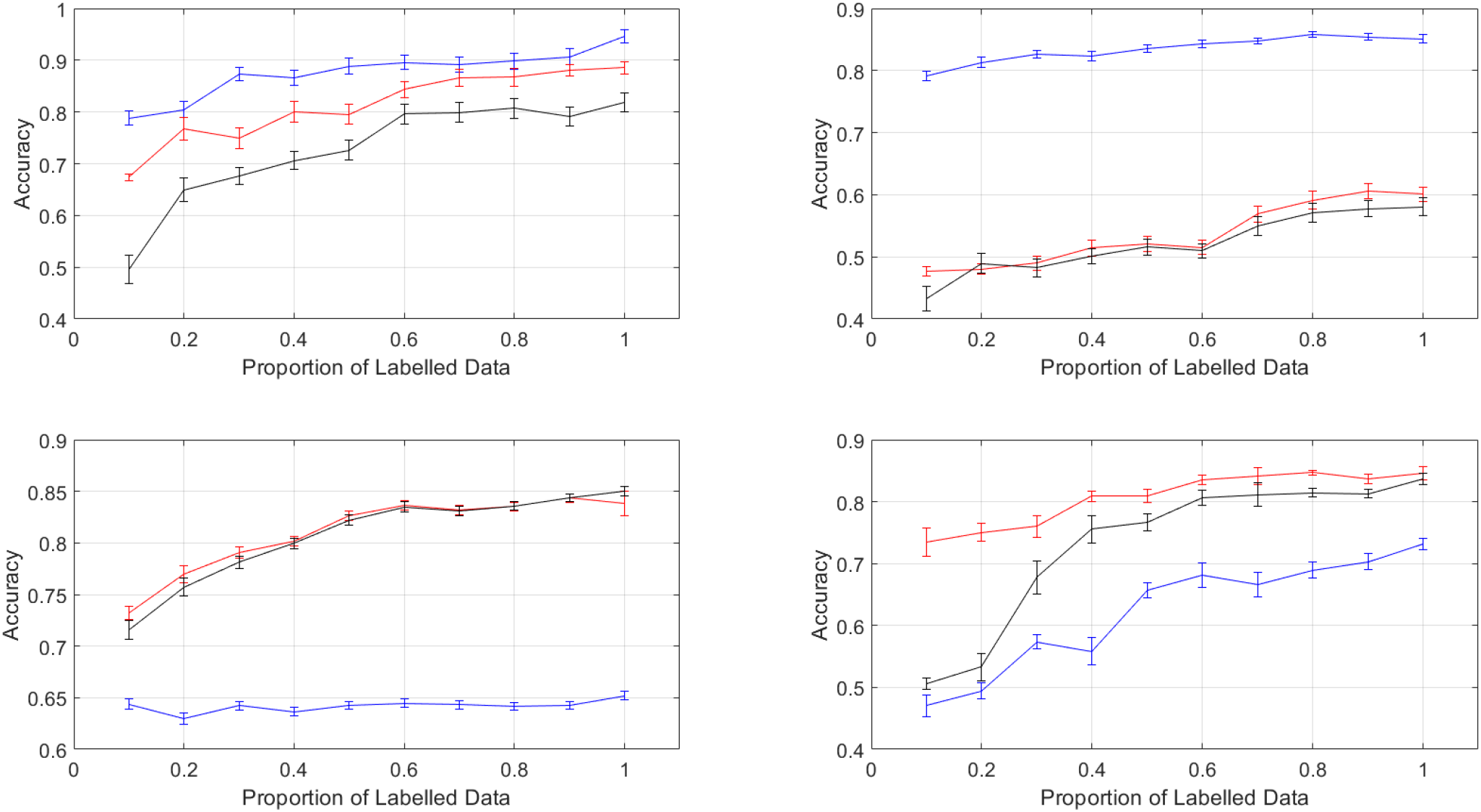
Semi-Supervised Learning. The graph shows test set accuracy versus the proportion of labelled training data for the Cluster-Based Memory Module (CMM) (blue curve), Online Softmax Regression (black curve) and Batch Softmax Regression (red curve) for the (top-left) Iris, (top-right) Ecoli, (bottom-left) Ionosphere and (bottom-right) MNIST data sets. The plots show the mean test-set accuracy, and standard error of the mean, over multiple random partitions of the data. CMM provides better performance for Iris and Ecoli, but worse for Ionosphere and MNIST.

### 3.4 Cluster Merging on Benchmark Data

Here we show the effect of cluster merging on the ML data sets. On each simulation run we randomly split the data into equal sized training and testing sets and assessed classifier accuracy using the weighted balanced accuracy on the test set. This was repeated for ten simulations each of which used 100% labelled data. As can be seen from table 2, merging naturally results in fewer clusters. This significantly improved accuracy on the MNIST and Ionosphere data, albeit from rather low baselins. Merging did not improve categorisation accuracy but did improve memory efficiency in that fewer clusters were required.

**Table 2:** Cluster Merging on Benchmark Data. Accuracy is the weighted balanced accuracy on the test set and the asterisk denotes a significant difference at p ≤ 0.05.

| Data Set | Number of Clusters |  | Accuracy |  |
| --- | --- | --- | --- | --- |
|  | CMM | CTM | CMM | CTM |
| Iris | 5 | 3 | 0.93 | 0.93 |
| Ecoli | 11 | 4 | 0.86 | 0.87 |
| Ionosphere | 37 | 12 | 0.65 | 0.71* |
| MNIST | 99 | 52 | 0.73 | 0.75* |

## 4 Discussion

This paper has proposed a Cluster-Based Inference framework for describing how the brain might capture and organise experience and how these representations can be efficiently transformed in the service of a specific task such as categorisation. We have shown how many of the attributes of memory-based cognition could be provided by such a system including those of recognition, causal attribution/inference, pattern completion, and generative replay. Within this we have proposed that the opposed goals of providing a system with strong pattern separation of learning episodes and strong generalisation across thoses episodes be instantiated by a Cluster-Based Memory Module (CMM) and a Cluster-Based Task Module (CTM), respectively. In a categorisation learning context, the CMM maximises the joint probability of stimul and labels whereas the CTM maximises the conditional probability of labels given stimuli.

### 4.1 Behavioural Data

Our simulation results on the behavioural task data find that cluster merging improves generalisation, at a cost of memorisation, but overall yields improved performance. A consequence of cluster consolidation is that model-based recognition scores for old items are considerably reduced for old cues. These simulation results therefore indicate that in this context forgetting is a natural consequence of optimising for future performance, a finding also reflected in other perspectives of memory function [56]. A second finding was that, after cluster consolidation, model-based recognition scores were considerably increased for new cues indicating a potential “false memory” effect [29]. Again, the explanation here is that this is in service of future (categorisation) performance.

The behavioral task could not be learnt accurately with a low precision stimulus encoding during online learning. Use of a low precision encoding typically resulted in a single cluster model without any category specificity. For good results, it was necessary to use a high stimulus encoding precision during online learning. This results in a large number of clusters. Cluster merging, driven by the goal of maximising the conditional probability of categories given stimuli, then results in fewer but larger clusters.

As the encoding scale approaches zero the number of clusters approaches the number of unique items in the training set. The model therefore takes on characteristics of an episodic memory, having a unique representation for each unique stimulus, operating similarly to a configural reinforcement learning model [16] which learns reward/decision probabilities for each unique cue configuration. This strong pattern separation domain is characteristic of the CMM.

The experimental findings from human behavioural data [47] also showed an improvement in generalisation. This was significantly non-zero within the Quiet Wakefulness (QW) group, and significantly greater for the QW than a control group. However, no reduction in performance for old cues (“memorisation”) was found for either the QW or control groups. The model predictions of better generalisation but worse memorisation are therefore only partially consistent with experimental findings. These findings are from an online experiment and future work will aim to replicate these in a lab-based setting as with other behavioural research in offline learning [14].

### 4.2 Benchmark Data

CMM provided better performance for the Iris and Ecoli data sets with improved performances available using even a small proportion of labelled data e.g. 10 per cent. As Softmax Regression is a linear classification method, these improved performances are likely due to nonlinearities in decision boundaries; CMM can exploit these but Softmax Regression can’t. CMM provides worse peformance for Ionosphere and MNIST data, with performance at chance level for Ionosphere. We don’t know why CMM fails for these data but one possibility is the increased data dimension (see *d* in Table 1). CMM produces a large number of clusters for both of these datasets (see Table 12) and having a large number of underlying parameters is not usually associated with good pattern recognition performance.

Cluster consolidation does reduce the number of clusters and therefore parameters and performance is significantly improved for CTM over CMM. However, the absolute levels of performance for these data are not comparable to what is achievable with other methods. We therefore conclude that neither CMM or CTM are useful as practical decision making algorithms even with medium dimension data (tens of dimensions). One possible reason for this is that cluster consolidation has been implemented using only a merging process, whereas standard model selection algorithms in machine learning often use a combination of merging, splitting and birth/death steps. A further alternative that is within the CTM framework, but not implemented in this paper, would be to implement CTMs using a Mixture of Experts type algorithm whereby within-cluster probabilities are not constant within cluster (as for the current CTM) but vary as a function of stimulus input. This could be implemented using Softmax Regression experts and will be investigated in a follow-up paper.

### 4.3 Model-based Neuroimaging

A key marker of cluster consolidation is that novel stimuli presented at test will produce higher recognition scores than if no consolidation took place. A model-based neuroimaging approach based on the behavioural task examined in this paper should be able to identify the neural correlates of these scores and therefore identify where in the brain cluster consolidation occurs. Given this experimental paradigm has two test epochs sandwiching the offline wake period, a regressor can be created that is the change in recognition score for each novel item (i.e. for cues not presented in training). Brain regions that are correlated with this change in recognition score will then be identified as the loci of cluster consolidation. This procedure would be similar to the fitting of RCMs [15, 45] and LCMs [1, 58] to participant’s decision time series which, in related paradigms, have shown recognition scores to be associated with activation of posterior hippocampus [15].

### 4.4 Neurobiology

Given that the spatial scale of grid and place cells increases as one proceeds anteriorly in the hippocampus [64] and that these cells also provide representations over non-spatial domains [12], it may be the case that merged models are instantiated in anterior hippocampus. An alternative perspective is that merged models may instead be instantiated in vmPFC or that this region is involved in the creation of CTMs. According to Gilboa and Marlatte [29], “… during formation of overlapping associations, the vmPFC exerts control that allows conflict resolution between competing associations in hippocampus and the creation of an integrated schematic representation”. Intriguingly, there is evidence that an intact vmPFC is necessary for a range false memory effects [29]. These false memories may be analagous to the inflated recognition scores for novel cues from the merged model (see above).

### 4.5 Model Evidence

The cluster consolidation algorithm we have investigated uses the variational free energy, an approximation to the log of the Bayesian model evidence, as a model selection criterion for Cluster Based Models. The number of clusters is determined by the model with the highest model evidence. Cluster-merging stops when the model evidence decreases.

This approach is similar to that of Heller and Ghahramani [35] who show that the model evidence can be used as a merging criterion, and be applied recursively in the context of hierarchical clustering. Use of the Bayesian model evidence has been proposed as a mechanism which the brain may use to adjudicate among its many and various prediction systems [21].

In related work, Smith et al. [62] use an active inference framework to solve simple concept learning problems. The approach is similar to that in the current paper in that the number of latent variables, or “slots”, can expand to accommodate learning of new concepts, but these slots can also be selectively removed, or increased in granularity (c.f. cluster consolidation), to aid generalisation. Slot updating is governed by the maximisation of model evidence.

More generally, in the statistics and machine learning literature model selection algorithms have been proposed that rely on cluster-splitting and cluster-pruning as well has cluster-merging operations [5]. However, as our goal in this paper was to operationalise a proposal from cognitive neuroscience [73] we have focussed solely on cluster merging. Cluster-pruning and splitting may be additional processes that are engaged, for example, during reward-based learning [60] or synaptic downscaling [44].

Fountas et al. [23] propose an active inference model of episodic memory with split/merge steps based on Jensen-Shannon (merge) or Kullback-Liebler (split) criteria. However, these criteria scale with the number of model parameters and must be individually optimised for each application. In contrast, differences in log model evidence are equivalent to log Bayes factors, and have a natural scale [39] that is independent of data or parameter dimension.

### 4.6 Latent Cause Models

Latent Cause Models (LCMs) [13, 26] of classical conditioning also employ an adaptive cluster-based approach but one which provides a joint density over sensory stimuli, contextual variables and rewards received. Here, the ‘causes” are cluster assignments. The probability of reward given stimulus and context can then be computed using pattern completion. Latent Cause Models (top of p. 284 in [54]) can exhibit one-shot learning e.g. in reversal learning paradigms by inferring that a cause or context has changed. They also explain (i) the ‘spontaneous recovery effect” in associative learning whereby when a neutral stimulus is no longer paired with a shock, the associated fear response quickly disappears but later spontaneously returns, (ii) the ‘gradual extinction effect’ – such that if the neutral stimulus is subsequently paired with gradually decreasing amounts of shock, extinction becomes more permanent. This is because, for (i) a new latent cause is created, whereas for (ii) the old latent cause is gradually adapted.

### 4.7 Mixture Models

Ghaharamni and Jordan [28] have shown how to use mixture models for supervised learning. In the learning stage the mixture model estimates the joint density over all inputs and outputs. In the testing phase, the conditional density of outputs given inputs can then be computed from the joint density learnt in training, using Bayes rule. This is identical to the ‘pattern completion’ described in this paper. They have described the implementation of this for mixture models of both real-valued and discrete variables using mixture components with both Gaussian and multinomial probability distributions. These are also known as ‘shared component’ mixture models, because clusters can be shared by different categories [38]. This is the same model that underlies cluster-based models for category learning. The only differences are that we have added task variables (although they are not made use of in the current paper) and have used variational inference to provide a model selection metric.

### 4.8 Deep Autoencoders in Neocortex

In this paper we have used a Variational Autoencoder (VAE) [40] to reduce the dimension of the MNIST character recognition data set. This fits in with an assumed neurobiological correspondence in which autoencoders provide reduced-dimension stimulus representations available in multimodal association cortex which then provide input to hippocampus. In related work, Hedayati et al [34] use a multiple-layer autoencoder model of the ventral visual stream as part of a visual working memory model. Here the autoencoder makes use of statistical redundancies within a single sensory modality, vision, to produce a low-dimensional code that can be projected to association cortex. Additionally. Ven et al. [67] use a deep autoencoder as part of a continual learning system – this is discussed further in the consolidation section below.

### 4.9 Shallow Autoencoders in Hippocampus

The hippocampal system may itself perform further dimensionality reduction internally [30], making use of the statistical redundancies among sensory modalities (e.g. correlations among visual, auditory, olfactory and emotional percepts). This can also be instantiated with autoencoders but perhaps with a shallower architecture than neocortex (with most implementations comprising a single hidden layer). Autoencoding is central to the Gluck-Myers hippocampal representation theory [30] and a recent model of this type shows how place cells can be learnt [6].

The training of autoencoders usually employs a batch learning algorithm and so is not online. This therefore makes them unrealistic as models the sort of “fast” hippocampal learning that is envisaged in complementary learning systems theory. It may be the case, however, that online learning is indeed possible, at least for autoencoders with a single hidden layer where nodes can be added in a nonparametric approach [75].

A variant of nonlinear autoencoding could be implemented in CMMs and CTMs by replacing the Gaussians with factor analysers [4]. The stimulus-facing part of the model would then be a nonparametric mixture of factor analysers (rather than a nonparametric mixture of Gaussians). Although the encoding is linear within each cluster, because there are multiple clusters the overall coding would be nonlinear (see [8] for an example of this type of approach in image coding).

### 4.10 Temporal Dynamics

A major drawback of our framework as a model of hippocampus is that it has no temporal structure. This may seem particulary odd as one of the main proposed functions of the hippocampal formation is as an episodic memory in which recall fidelity, for example, depends on order effects and elapsed time since encoding [37]. Probabilistic models of memory also focus on its sequential nature [63, 23]. The recently proposed Tolman-Eichenbaum Model [69] embeds temporal dynamics within an autoencoder framework. This employs both a hierarchical model, with two layers of latent variables, and can accomodate nonlinear temporal structure in each of those layers. The top level of the hierarchy describes variables *g_t_* whose activity evolves according to a transition density *p*(*g*_*t*+1_|*g_t_, a_t_*) where *a_t_* is an agent’s action at time *t*. The lower level describes variables *z_t_* whose activity evolves as 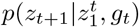 where 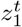 denotes the activity up to time *t*. Finally, observations *u_t_* are given by *p*(*u_t_|z_t_*). This nonlinear hierarchical dynamical model explains diverse phenomona from the development of grid-cell (*g_t_*) and place-cell (*z_t_*) like representations in spatial learning contexts to relational inference in more abstract contexts. Moreover, a fast Hebbian learning mechanism can be used for one-shot learning of place cell dynamics. This fast hippocampal learning is contrasted with the slow learning of relational structure in neocortex and is broadly consistent with CLS.

### 4.11 Systems Consolidation

The focus of this paper has been on Cluster Consolidation, but here we briefly review relevant work in the area of Systems Consolidation.

Ven et al. [67] use a deep autoencoder architecture as part of a continual learning system. They replaced the standard Gaussian prior over latent variables in a VAE with a *Gaussian distribution for each category* (see Fig 7 in that paper). This allowed category-specific examples to be generated as part of a generative replay mechanism and was a key element in providing state-of-the-art performance. The representations replayed were those in the highest (deepest) level of the autoencoder. Barry and Love [3] took a similar approach but employed a more neurobiologically realistic deep learning model of neocortex with five blocks of convolutional layers that could be associated with specific human brain regions. The system had also been trained on a larger set of categories (1000 versus 10). Here, they examined the benefit of replay for learning to categorise exemplars from 10 novel categories, as a function of which representations were replayed to cortex and found that benefit was maximal for representations in later versus earlier layers (putative lateral occipital cortex versus early visual cortex). Similar to Ven et al., replay was instantiated by fitting a single Gaussian distribution to distributions experienced during ‘awake’ experience, separately for each category, and samples were generated from these distributions during ‘sleep’.

Unlike both of these approaches, the probability of observed representations (stimuli) in our model is not directly conditioned on category labels. That is, we don’t for example have a separate Gaussian for each category of data. Indeed our model is richer, in that clusters can be shared among categories (this also constrains the overall complexity of the model). By instantiating the category (see ‘Generative Replay’ in section 2.1), however, we can activate the relevant clusters and then sample from those activated Gaussians. This produces replay data which is effectively from a *Gaussian-Mixture model for each category*. In principle, this should be superior to Gaussian approaches [67, 3] but we shall see how effective it is in future work.

We hypothesise that Barry and Love found that replay was beneficial for later layers because the distribution of activations in those layers was assumed Gaussian (given category labels), whereas category-conditional distributions of activations in earlier layers are likely to be more complex and multimodal. However, the earlier one goes in neocortex the more difficult it will be to specify an accurate probabilistic model of those distributions (in the limit of going back to primary sensory cortex the model would need to have a similar complexity to that of a neocortical autoencoder or deep learning model), so it would make sense for replay to be focused on higher-level activations. Moreover, it is unlikely that layers in early sensory cortex remain plastic in the adult human brain, so replay of those activations would serve no purpose.

Sun et al. [65] propose a theory of systems consolidation in which hippocampal-cortical interactions serve to optimise generalisations that guide future adaptive behaviour. In their formalism, a hippocampal notebook (instantiated as a sparse Hopfield net) encodes a multivariate pattern that is comprised of both independent variables (e.g. stimulus vector) and a dependent variable (e.g. category label), whereas the cortex encodes a (linear) mapping from independent to dependent variable. In statistical terms, this corresponds to our distinction between CMMs (hippocampal cluster-based memory module) and NTMs (cortical neural-net based task module). They show that if the mapping to be learnt is very noisy then hippocampal pattern-replay can, at some point, become detrimental to cortical learning. This leads to the proposal that replay must be regulated to promote generalisation. This principle, and their computer simulations, explain various empirical findings regarding the time course of consolidation and retrograde amnesia. They show that multiple interacting memory systems can outperform just one and that the predictability of experiences should determine when and where memories reside.

They do not specify what process may govern the regulation of replay. One suggestion, however, is that “memory elements reflecting predictable relationships could be replayed together”. This would fit with the idea that samples from merged cluster based models (CTMs) would be produced during generative replay, rather than from unmerged models.

### 4.12 Cost Functions

The central idea of this paper is that a probabilistic model of episodic memory should maximise the joint probability of multi-sensory inputs and e.g. category labels (and any other attended information available at the time of encoding), whereas a memory-based system that is optimised for task performance, such as categorisation, should maximise the probability of category labels given sensory input, and the creation of the latter through a model-based consolidation process might explain a number of phenomena in memory research e.g. forgetting, false memories, and why some long-term memories remain dependent on hippocampus. This central idea may be correct even though the instantiation of it with cluster-based models may be of more limited value. We also wish to emphasise that we envisage the above as sitting within an overarching process of systems consolidation that likely operates at a longer time scale due to differences in the nature of the representations – highly distributed for neocortex versus more local in hippocampus.

## A Log Joint

To derive the variational learning algorithm we first need to write down the “log-joint”, that is, the log of the joint probability of data and latent variables. This is given by

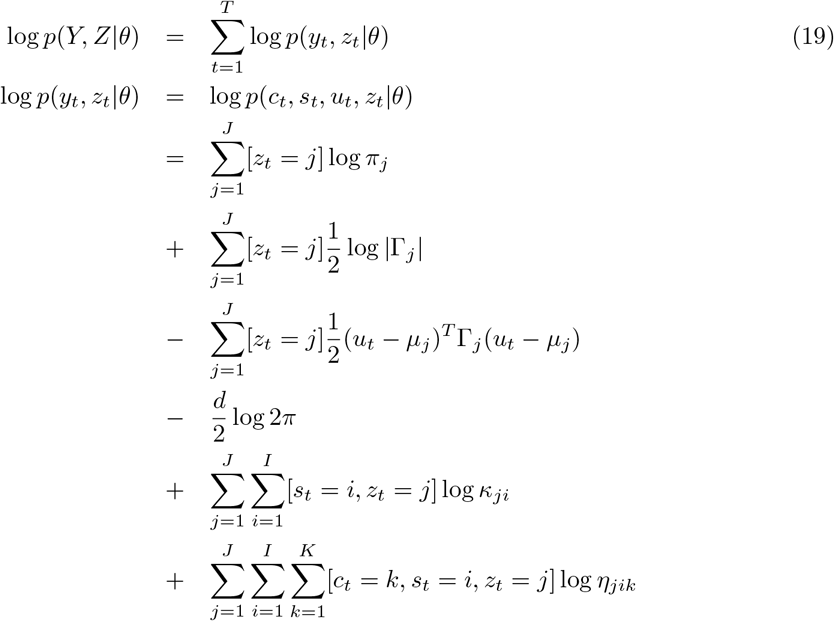

## B E- and M-Steps

For the likelihoods and priors in the Cluster-Based Model we don’t need to assume a particular factorization or functional form for the factors. These forms fall out of what is referred to as a free-form optimisation of the variational objective function [2, 4] (this is to be contrasted with fixed-form approaches which are necessary when the likelihoods and/or priors are not in the exponential family e.g. [25]). Given the additive structure of the log-joint over t, the approximate posterior over latent variables is given by (see [2] for generic procedure)

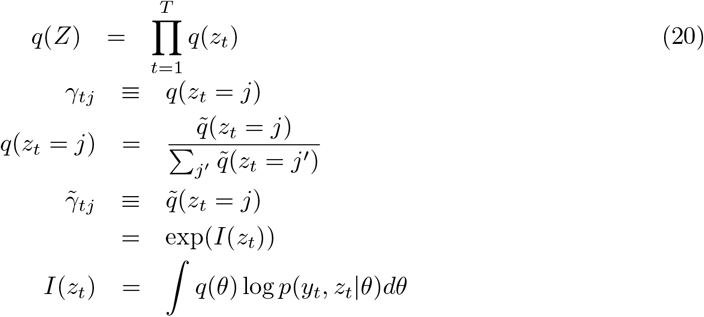

where *γ_tj_* is the probability that cluster *j* is responsible for category, task and cue data at time *t*. We also refer to the un-normalised probabilities, 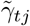, as the ‘cluster activations’. By substituting the log-joint from Eq. 19 into equation 20 and integrating over *q*(*θ*) we can derive the E-Step equation described below.

The approximate posterior over parameters is given by [2]

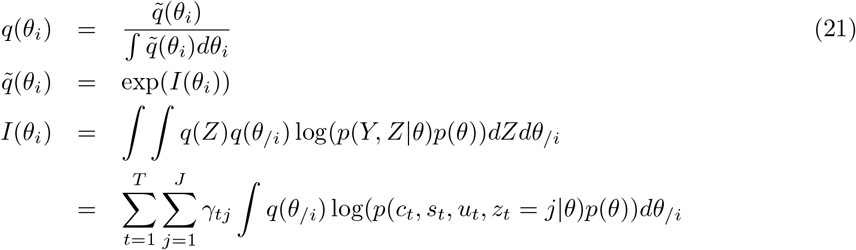

where *θ_i_* refers to parameters in the ith group, and *θ/i* to all other parameters. In the Cluster-Based Model the parameters are grouped according to the structure of the prior (see Eq. 9). By substituting the log-joint from Eq. 19 and log of the prior from Eq. 9 into equation 21, and integrating over *q*(*θ/i*) we can derive the M-Step equations described below.

### B.1 E-step

The E-step consists of updating the cluster posterior as follows. First we compute a number of preliminary quantities

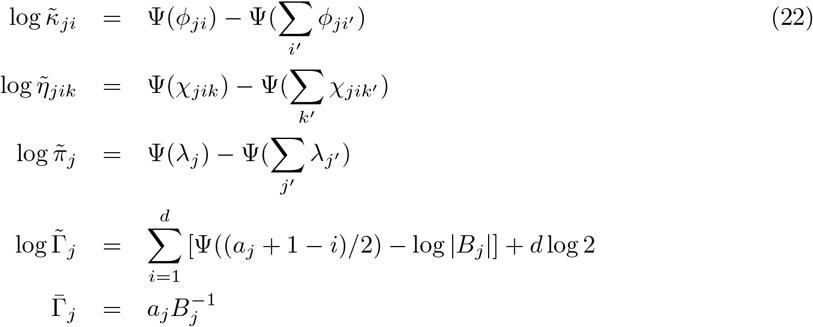

where Ψ() is the digamma function. We then compute the ‘cluster activations’ (unnormalised cluster posteriors)

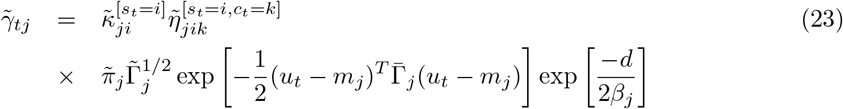

Equation 23 shows that the cluster activations are a product of incoming signals with a term for each data type (here – task, category, and cue). As we discuss in the main text, this mathematical form reflects the fact that experiences are multidimensional, and may correspond to conjunctive representations in hippocampus [70]. The cluster posteriors are then computed in a final normalisation step

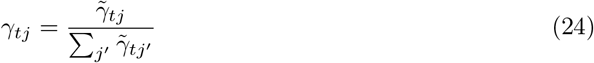

### B.2 M-step

First of all, we define the following variables

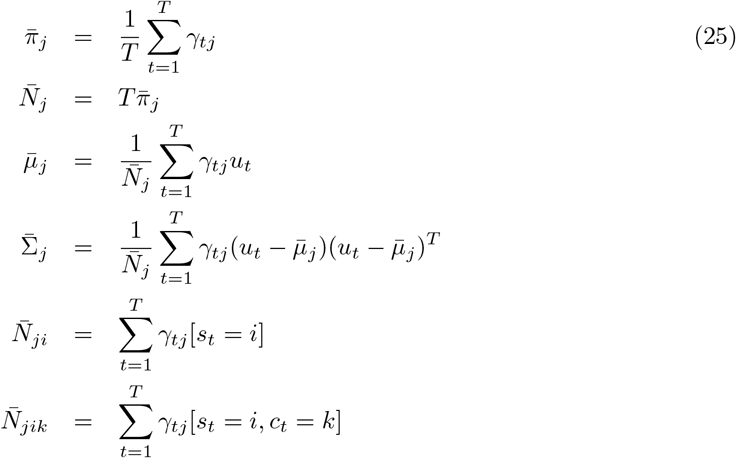

where 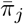 is the proportion of data assigned to cluster *j* and 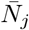 is the number of data points in cluster *j*. We then update model parameters as follows. For the Dirichlet distributions we add data counts to prior counts

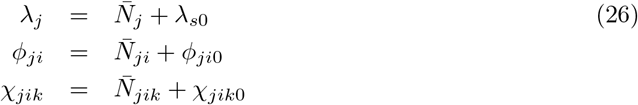

For the means, we have

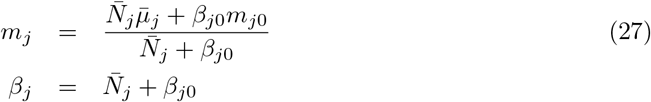

For the noise precision matrix we have

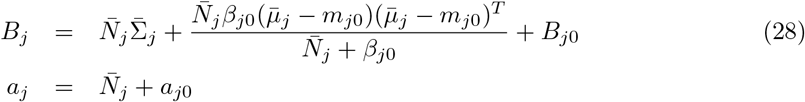

## C Model Evidence

The negative variational free energy, *F*, provides a lower-bound approximation to the model evidence

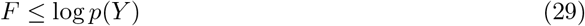

for a model of data *Y*. For models with parameters *θ* and latent variables Z this can be expressed as

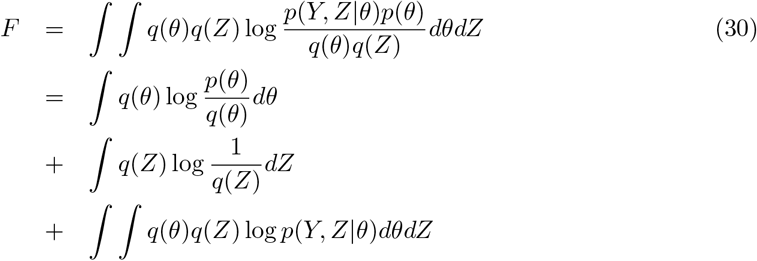

The quantity *F* is also known as the Evidence Lower Bound (ELBO) [40] and is the same quantity that is central to the Free Energy Principle of systems neuroscience [24].

### C.1 Joint Density

For the CMM we have data *Y* = {*C, S, U*} and model parameters *θ* = {*η, κ,μ, Γ, π*}. We can compute a lower-bound approximation to the evidence of the CMM

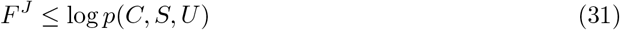

Substituting the model densities into Eq. 30 gives

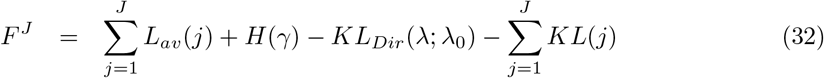

where

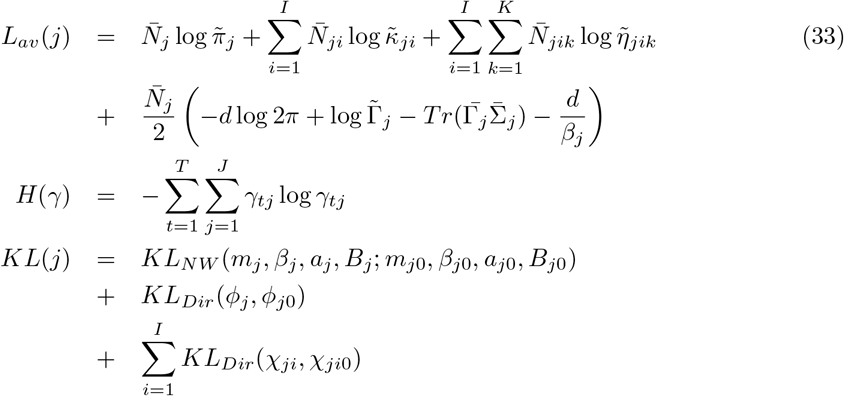

where *H*(*γ*) is the entropy of the cluster responsibilities (all else being equal, clusterings with higher entropy result in higher model evidence) and *KL* denotes the Kullback-Leibler Divergences which are defined in [53]. *KL_NW_* and *KL_Dir_* denote the divergences over Normal-Wishart and Dirichlet distributions respectively.

### C.2 Conditional Density

In the CTM, or ‘categorisation model’, the stimuli *U* are not treated as random variables generated by the model. They are treated as known inputs. Additionally, at the time of making categorisation decisions, the task variable is also known. Our probabilistic model is therefore over category variables only

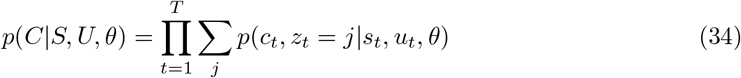

We can compute a lower-bound approximation to the evidence of the CTM

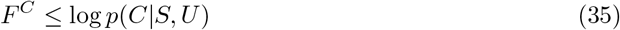

Our data are now *Y* = {*C*}, the parameters are *θ* = {*η, κ, μ, Γ*}, and an expression for *F^C^* can be determined by substituting the densities from the CTM into Eq. 30. One density we need is the joint density over category labels and latent variables, *C* and *Z*, which can be expressed as

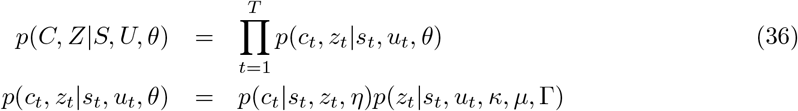

The substitution then produces

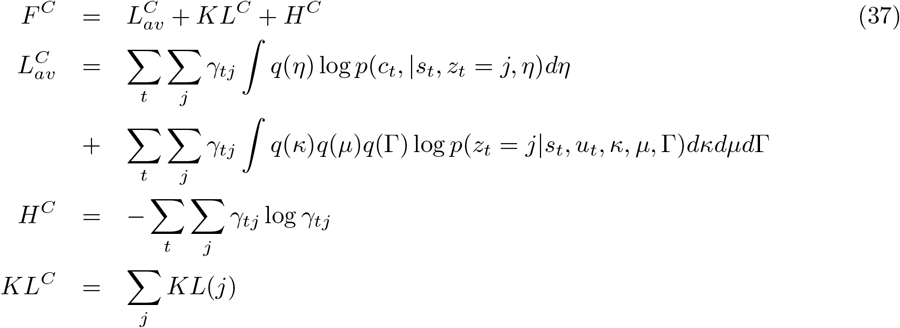

where *KL*(*j*) is exactly the same as for the CMM. The integral in the second line of 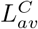 is equal to log 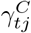 where

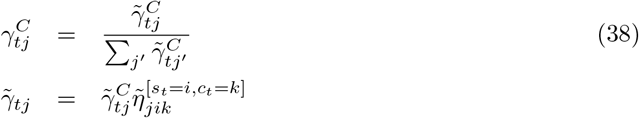

The quantity 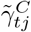 is therefore given by Eq 23 but with 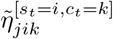 removed. We can therefore write

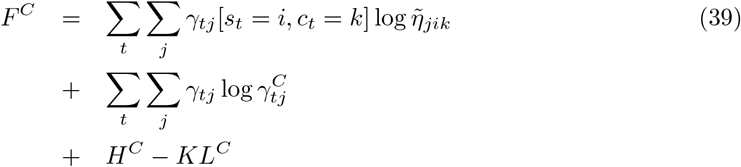

which can be written more compactly as

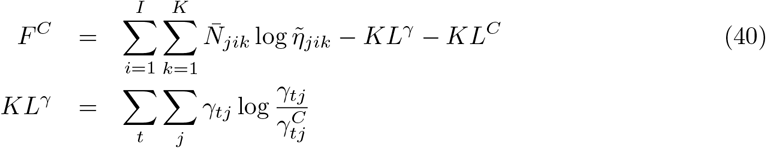

where the first term in *F^C^* reports correct classification likelihood, *KL^γ^* penalises CTMs with different cluster assignments to CMMs, and *KL^C^* penalises models with many clusters.

## D Recognition Scores

The recognition scores which are used to compute the values in Fig 11, and can be used to create regressors for model-based neuroimaging are computed as follows. First, we note that the cluster activations, 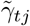 in Eq 23 are equivalent to the joint density *p*(*u_t_, c_t_* = *k,s_t_* = *i,z_t_* = *j*). The recognition scores are then given by integrating this quantity over category, task, and cluster. That is,

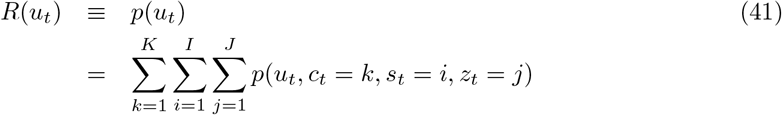

This is analagous to the example in Eq 4 except, in addition to *u* our the CMM additionally models the density over category and task labels. The recognition score *R*(*u_t_*) is the probability density at *u_t_* as estimated by the CMM. Recognition scores can be similarly computed for the CTM.

## E Category Probabilities

The category probabilities which are used for classification are computed as follows. First, we compute the joint density over stimuli and categories by integrating over the full joint density (see previous section). That is

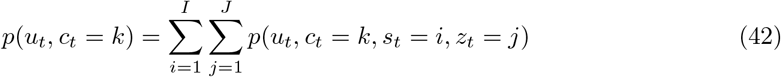

Category probabilities are then computed as

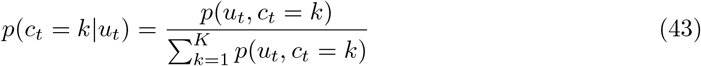

This is analagous to pattern completion shown in Eq 7 except now we are inferring a category variable from stimulus variables, rather than inferring one stimulus variable from another. This and the previous section are two examples of the general principle that all probability distributions of interest can be derived from the joint density.

